# Dopamine waves as a mechanism for spatiotemporal credit assignment

**DOI:** 10.1101/729640

**Authors:** Arif A. Hamid, Michael J. Frank, Christopher I. Moore

## Abstract

Significant evidence supports the view that dopamine shapes reward-learning by encoding prediction errors. However, it is unknown whether dopamine decision-signals are tailored to the functional specialization of target regions. Here, we report a novel set of wave-like spatiotemporal activity-patterns in dopamine axons across the dorsal striatum. These waves switch between different activational motifs and organize dopamine transients into localized clusters within functionally related striatal subregions. These specific motifs are associated with distinct task contexts: At reward delivery, dopamine signals rapidly resynchronize into propagating waves with opponent directions depending on instrumental task contingencies. Moreover, dopamine dynamics during reward pursuit signal the extent to which mice have instrumental control and interact with reward waves to predict future behavioral adjustments. Our results are consistent with a computational architecture in which striatal dopamine signals are sculpted by inference about instrumental controllability and provide evidence for a spatiotemporally “vectorized” role of dopamine in credit assignment.

---

Dopamine supports reward learning and motivational activation, but details about what decision variables are encoded, and how they are delivered to postsynaptic targets, continue to be refined^(Berridge, 2007; Schultz, 2016; Berke, 2018)^. The dopamine-reward prediction error (RPE) hypothesis emphasizes that dopamine conveys deviations from reward expectation in reinforcement learning (RL) theory^(Schultz et al., 1997)^. This formulation generally treats dopamine as a “global” spatio-temporally uniform signal, a view based on two key findings. First, extensively divergent dopamine axons^(Matsuda et al., 2009; Prensa and Parent, 2001)^ provide an architecture for broadcast-like communication. Second, dopamine cell spikes measured in the midbrain are highly synchronized^(Hyland et al., 2002; Li et al., 2011)^, putatively implementing a redundant population code^(Joshua et al., 2009; Kim et al., 2012; Mohebi et al., 2019)^ for RPEs^(Eshel et al., 2016)^. These observations form the basis for an influential view^(Glimcher, 2011; Schultz, 1998)^ of what dopamine communicates and how it is delivered: scalar RPEs that are uniformly delivered to all recipient subregions. The notion of uniform encoding also extends to alternative accounts for dopamine’s role in motivation^(Berridge, 2007)^ by relaying scalar value signals^(Hamid et al., 2016)^.

A key limitation of this global view is that scalar (or, spatio-temporally uniform) decision variables are neither computationally advantageous, nor reflected in forebrain dopamine dynamics. Postsynaptic striatal subregions are functionally specialized^(Graybiel, 2008; Haber, 2003)^, receiving distinct cortical and thalamic afferents^(Hintiryan et al., 2016; Hunnicutt et al., 2016)^, and express unique compliments of biomarkers^(Riedel et al., 2002)^. Accordingly, rewards^(Brown et al., 2011)^, their motivated pursuit^(Hamid et al., 2016; Shnitko and Robinson, 2015)^ and predictive stimuli^(Menegas et al., 2017)^ produce vastly different dopamine time courses across the dorsal-ventral and medial-lateral axes of the striatum. While these observations indicate regional heterogeneity, the extent to which dopamine inputs reflect the computational requirements of postsynaptic areas remains elusive. For example, there is some theoretical motivation^(Doya et al., 2002; Frank and Badre, 2011)^ and empirical support^(Badre and Frank, 2011; Gershman et al., 2009)^ for delivery of vector-valued RPEs that depend on a target region’s computational specialty. Nonetheless, we currently lack a clear understanding of organizing principles for striatal dopamine activity, and what normative computational functions may be served by such heterogeneity.

## Related striatal subregions get correlated dopamine input

We set out to characterize the spatio-temporal organizational rules of dopamine activity across the dorsal striatum. Standard methods for dopamine measurement typically survey small territories (10s – 100s of micrometers) and are ill-suited to probing large-scale organization. To overcome these limitations, we injected *cre*-dependent GCAMP6f into the midbrain of DAT-cre mice, and imaged dopamine axons through a 7mm^2^ chronic imaging window over the dorsal striatum (DS)^(Howe and Dombeck, 2016)^ (**Fig. 1a**). This approach provided optical access to 60-80% of the dorsal surface of the mouse striatum, with a view of dorsomedial (DMS), dorsolateral (DLS) and partial access to the posterior-tail (TS) region of the striatum (**Fig. 1b**). We imaged the activity of dopamine axons at multiple levels of resolution with one or two photon microscopy.

**Figure 1:**
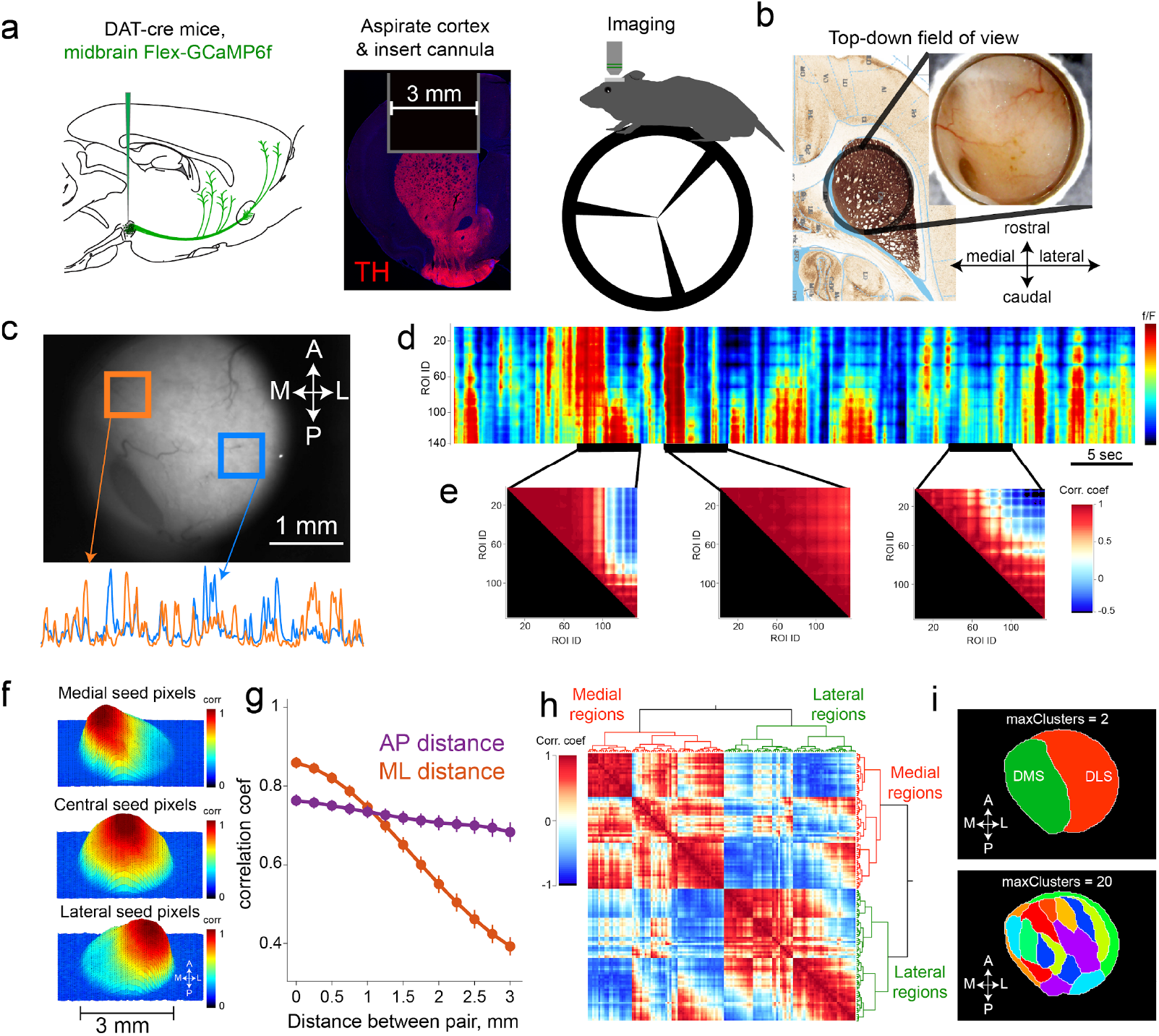
Dorsal striatal dopamine activity is spatiotemporally asynchronous and clusters into contiguous territories. **a**, Schematic of methods for imaging dopamine axons over dorsal striatum. GCaMP6 was injected into midbrain of DAT-cre mice. Cortex overlying dorsal striatum was aspirated, together with insertion of 3 mm diameter imaging cannula, and activity was imaged in head-restrained mice. **b**, Top-down field of view. **c**, Average delta f/F from two regions (top) that exhibit decorrelated activity (bottom). **d**, Activity of several ROIs from the same session as **c**, time series are sorted such that medial areas are top ROIs, and lateral regions are represented at the bottom. **e**, Correlation matrix across ROIs for different 5sec epochs (highlighted in bottom of **d**), showing patterns of correlations that evolve in time. For example, middle shows global correlation, whereas left and right panels show instances of antagonistic activity patterns in top and bottom set of ROIs. **f**, Results from spatial correlation from seed pixels, evaluating the Pearson’s correlation of with all other regions. *Top* panel shows medial seed pixels that are highly correlated with nearby regions and show graded decrease in correlation for distant regions. Same analysis was repeated for a set of pixels in central striatum (*middle*) and lateral seed pixels (*bottom*). **g**, Quantification of sessions-wide correlation between each pair of pixels as a function of distance, separated by medio-lateral (orange) and anterio-posterior distances. (n= 8 mice, p<0.001 wilcoxon signed-rank test for difference of ML vs AP slopes). **h**, Paiwise correlation matrix using hierarchical clustering summarizes similarity of dopamine activity. **i**, *Top*, anatomical projection of pixels that share similarity at the highest cluster limit of two outlining medial and lateral subregions of the dorsal striatum. Increasing the cluster threshold to 20 (*bottom*) revealed smaller, but anatomically contiguous regions of the striatum.

Using a head-fixed preparation, we began by focusing on spontaneous activation of dopamine axons in mice resting on a wheel in a dark chamber without external stimuli. To first test if dopamine axons were globally activated, we compared fluorescence signals in DS regions-of-interest (ROIs) (**Fig. 1c**). While ROIs were sometimes globally synchronized^(Howe and Dombeck, 2016)^, we observed decorrelated patterns across striatal subregions that evolved across time (**Fig. 1c,d,e**). These patterns of activation were observed across multiple anatomical scales (see **Supplementary Fig. 1** for micron-scale organization), indicating that dopamine afferents can become recruited asynchronously.

To examine how activity is spatially coordinated, we computed the Pearson’s correlation between pixels’ fluorescence as a function of anatomical distance. Dopamine axons showed strong local correlations that gradually decreased with distance (**Fig. 1f,g**), comparable to the organization of striatal spiny-neuron activity^(Klaus et al., 2017)^. Strikingly however, this distance-dependent falloff was selective to the medio-lateral axis, and was not present on the anterio-posterior axis (**Fig. 1g**), suggesting an organization rule that promoted selective mediolateral decorrelation.

To further examine the topographical organization of dopamine signals, we leveraged standard cluster analyses (**Fig. 1h**). In every dataset (n = 31 sessions, 8 mice), the highest cluster threshold identified two contiguous subregions outlining well-established^(Balleine et al., 2007; Yin and Knowlton, 2006)^ DS subregions; medial (DMS) and lateral (DLS) striatum (**Fig. 1i top**). Further increasing cluster limits progressively (**Supplementary Fig. 2**) revealed smaller subdomains of DS (**Fig. 1i bottom**), resembling striatal sub-clusters previously identified based on glutamatergic input patterns^(Hunnicutt et al., 2016)^. These areas had similar clustering patterns across days and animals (**Supplementary Fig. 3**), with 25-30 optimal clusters identified in our field of view (**Supplementary Fig. 4**). Shuffling the pixelwise temporal or spatial indices produced random clusters (**Supplementary Fig. 4**), indicating a critical dependence on the underlying spatio-temporal activity pattern. Together, these results provide evidence for regional coordination of dopamine transmission and provided an initial basis for evaluating whether these signals are modulated by the underlying subregion’s computational specialty.

## Wave-like patterns coordinate dopamine activity

What spatiotemporal patterns produce systematically decorrelated dopamine signals? We noticed that full-field fluorescence exhibited complex but spatially and temporally continuous trajectories throughout the striatum, similar to travelling waves described in other cortical and subcortical brain regions^(Grinvald et al., 1994; Lubenov and Siapas, 2009; Mohajerani et al., 2013; Muller et al., 2018)^ (**Fig. 2a,b, Supplementary Video 1**). To quantify these complex trajectories, we used optic flow algorithms^(Afrashteh et al., 2017)^ to compute frame-by-frame flow fields (see methods for details; **Supplementary Video 2**).

**Figure 2:**
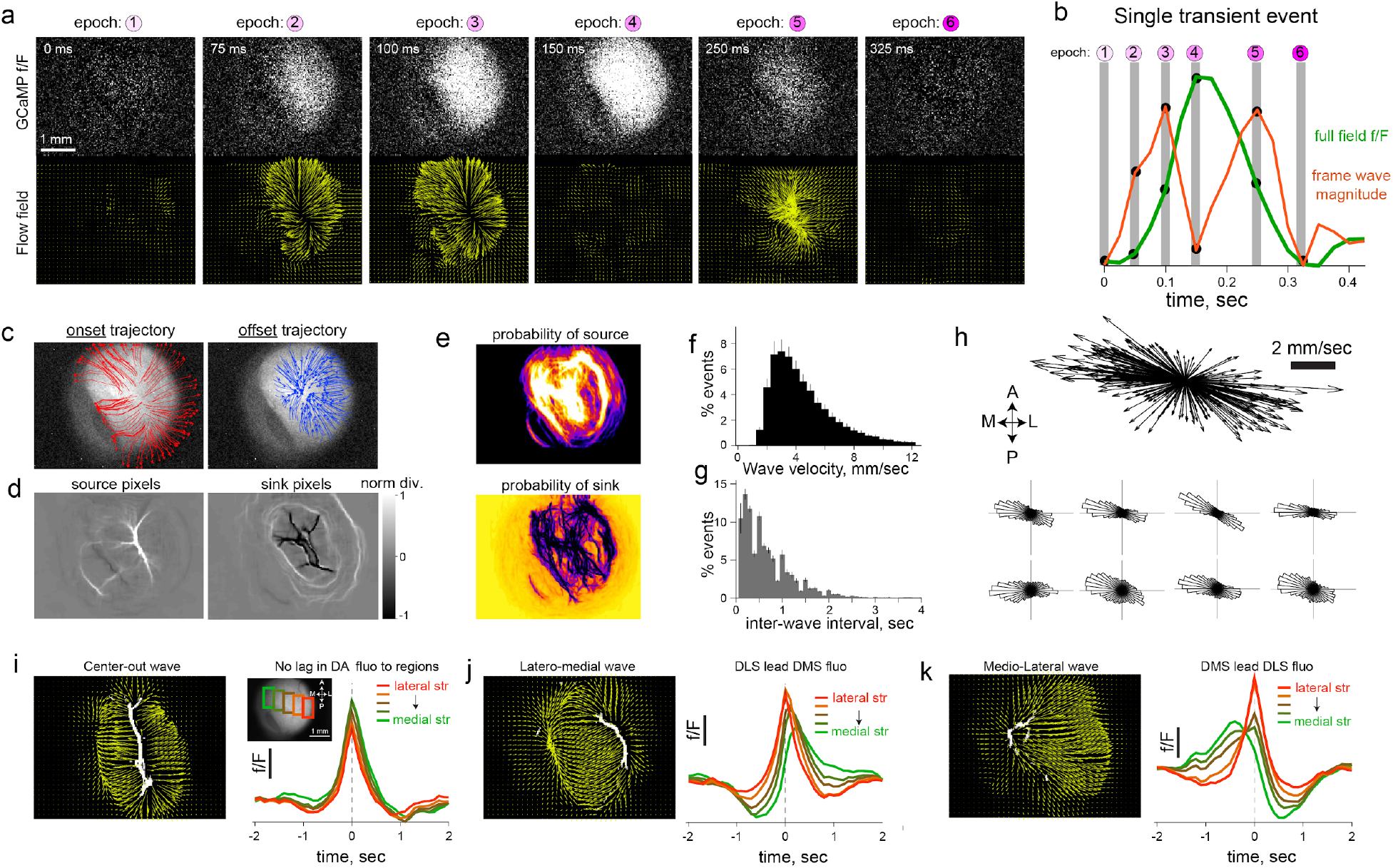
Wave-like spatiotemporally continuous sequences of dopamine-axon activation switch between motifs. **a**, *Top* row shows individual frames for different epochs of a transient as dopamine axon activity emerges and extinguishes in DS. *Bottom* row displays the corresponding flow vector fields computed for each pixel. Notice the divergence of vector fields during the rise phase of fluorescence, and convergent vectors during fall phase. **b**, Average fluorescence (green) across the entire field of view lasting ∼300ms sampled at 40Hz, and corresponding, flow magnitude in the fluorescence signal (red). **c**, Flow trajectory of fluorescence for 5 frames during the onset (*left,* red lines), or offset (*left,* blue lines) phase of the wave from **b**. Each line shows the pattern of flow from individually seeded pixels. **d**, Heatmap quantifying how divergent the vector fields are at each pixel during onset or offset of activity (left and right respectively). A positive value indicates that diverging pattern of flow at each pixel indicating that fluorescence is entering the striatum from those locations. Conversely, negative values are sink regions with converging flow vectors. **e**, Peak-normalized projection of the flow vector divergence for the onset (*top*) or offset (*bottom*) of all transients in one session (n=1516 events). Note that a repeated configuration of pixels serve as sinks and sources. **f**, Distribution of propagation velocity (n=8 mice, 1625 +/-213 events per mouse). Error bar denotes SEM.**g**, Distribution of interwave intervals for the same data **f**. **h**, *Top,* quiver plot summarizing the direction and magnitude of waves in a single session, and distribution of angle of wave propagation for each animal, *bottom* (n=8 mice, all p < 0.001, Omnibus test for angular uniformity). **i**, *Left,* vector fields (yellow) superimposed onto source pixels (white) for waves that are sourced at the midline and propagate bidirectionally outward. *Right*, corresponding fluorescence time course in ROIs on a medio-lateral gradient of the striatum (*inset*). **j,** Same format as **i**, for lateral source and medial flow or, **k**, medial source and laterally flowing wave.

The onset of activity in GCaMP fluorescence originated from clustered “source” locations, and rapidly migrated to other regions (**Fig. 2c,d, left**). By contrast, activity terminated as a result of flow toward “sink” locations (**Fig. 2c,d, right**). A repeated configuration of pixels had a high probability of serving as sinks and sources (**Fig. 2e, Supplementary Fig. 5**), indicating that local rules may dictate the initiation and termination of dopamine activity.

Dopamine waves entered the dorsal striatum with exponentially decaying inter-wave-intervals (**Fig. 2g**) and propagate with a range of velocities (median = 3.8 mm/s, interquartile range = 2.5, **Fig. 2f**). The overall direction of flow is bimodally distributed, with a biased medial-lateral propagation axis (**Fig. 2h**, all p < 0.001, Omnibus test for angular uniformity).

We next sought to determine if the collection of complex trajectories were made up of simpler, repeated sequences that may influence the time course of dopamine arriving at different parts of the striatum. Indeed, the combination of initiation loci and flow direction gave rise to motif waves that were scaled by propagation velocity and extent of striatum covered. We focused our attention on three motifs that produced most of the dopamine transients (**Supplementary Fig. 5**).

First, source pixels clustered at the juncture of DMS and DLS (**Fig. 2i**) initiated dopamine activity that rapidly spread bilaterally outward (Type-1*, “Center-Out” or CO wave,* **Fig. 2i, left**). These waves radiate across the striatum with the fastest velocities, arriving at all subregions with almost zero lag (**Fig. 2i, right**). Second, source pixels in lateral DLS initiated a wave that propagates medially (Type-2*, “latero-medial” or LM wave*). LM waves advanced across the striatum relatively slowly and delivered dopamine transients to DMS that were delayed relative to DLS in proportion to propagation speed (**Fig. 2j right**). Third, a medially sourced wave propagated laterally (Type-3*, “medio-lateral” or ML wave*, **Fig. 2k, left**), terminating in DLS. ML waves activate dopamine axons in the medial striatum first and recruit lateral regions with substantial delay (**Fig. 2k, right**). Together, these results demonstrate that wave-like patterns are a fundamental organizational principle of dopamine axonal activity, prescribing how activity initiates, propagates and terminates across DS.

## Rewards evoke directional dopamine waves

What is the functional role of dopamine waves in adaptive behavior? We set out to determine the computational significance of wave-like trajectories in the context of the well-studied role of striatal dopamine in instrumental behavior. The dorsal striatum exhibits graded behavioral specialty, with the DMS orchestrating goal-directed behaviors involving action-outcome contingencies, and the DLS implicated in stimulus-response behaviors^(Balleine et al., 2007; Corbit and Janak, 2010; Yin and Knowlton, 2006)^. Inactivation or manipulation of dopamine in DMS degrades goal-directed planning and action due to an inability to learn whether rewards are under instrumental control^(Balleine and O’doherty, 2010; Wunderlich et al., 2012)^.

We thus designed two operant tasks intended to manipulate action-outcome contingencies, and asked whether dopamine dynamics carry information about the degree of instrumental controllability (**Fig. 3a,b,c**). First, in an ‘instrumental’ task, rewards were contingent on mice running on a wheel to traverse linearized distance, with the progress to reward indicated by an auditory tone that escalated in frequency (**Fig. 3b,d**). On each trial, the distance that was needed to run for tone transitions (and ultimately, reward) was randomly selected from a uniform distribution (50-150 cm, **Fig. 3c, left**). Thus, while the mouse was in control of tone transitions, the specific contingencies varied across trials. A second ‘pavlovian’ task was administered in separate sessions. The task structure was identical except tone-transitions occurred after fixed durations within a trial (randomly drawn, 4-8 sec, **Fig. 3c right**), and progress to reward was unrelated to running. Trained mice exhibited anticipatory lick trajectories that increased with ascending tone frequency in both tasks (**Fig. 3e,f)**, indicating that mice used these tones to update their online judgment of progress to reward. Analysis of run bouts (**Supplementary Fig. 6**) revealed that mice invested goal-directed effort to receive rewards selectively in the instrumental task.

**Figure 3:**
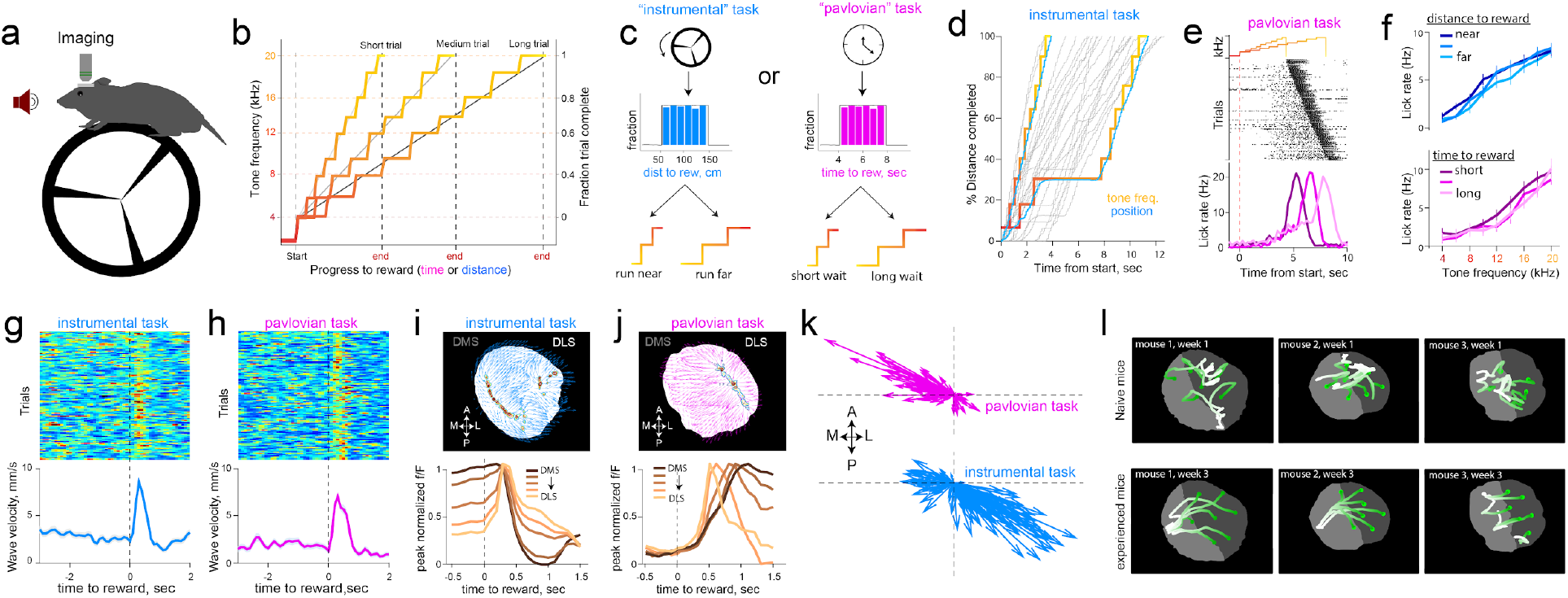
Reward delivery promotes directional waves that depend on instrumental requirement of task. **a**, Schematic of test chamber. **b**, Changes in tone frequency for short, medium and long trials tiling fraction of trial complete. **c**, In the instrumental task (*left*), tone transitions that escalate in frequency are linked to rotation of the wheel. The total distance to travel on each trial is drawn from a uniform distribution of 50-150 centimeters. In pavlovian task (*right*), the passage of time escalated tones, and the duration to wait was also drawn form a uniform distribution of 4-8 seconds. **d**, Example trials in the instrumental task. When the mouse traverses linearized distance rapidly, the tones escalate quickly, but if the mouse pauses running, tones do not escalate but still signal the fraction of distance completed. **e**, Example licking behavior in pavlovian session, sorted by delay to reward. Mice increase lickrate in anticipation of reward. **f**, These anticipatory licks were not influenced by distance to run, or duration to wait, but increased in proportion to progress to reward signaled by tones (two-way ANOVA effect of tones F(8,683) = 3.32), p = 0.001 and effect of 4 duration bins F(3,683)=0.48, p = 0.7 in pavlovian sessions. For instrumental sessions, effect of tones F(8,359) = 8.41, p<0.001 and influence of four distance bins F(3,359) = 0.13, p=0.9). **g**, Alignment of trial-by-trial reward wave velocity across the striatum. Rewards consistently resynchronized dopamine axons into waves in both the instrumental task (n=123 trials), and the pavlovian task **h** (111 trials)**. i,** *Top,* example flow vectors (blue arrows) and source locations (contour plot representing source regions) across pixels for a single rewarded trial. *Bottom,* peak-normalized fluorescence time course across trials produced by mediolateral waves on the medio-lateral gradient of the striatum. **j,** Same format as **i** for pavlovian session. **k,** Flow vectors for reward epoch (0-1s post reward across all pixels) for each trial in pavlovian and instrumental sessions shown in **g** and **h**. Each arrow shows the mean flow direction across striatum averaged across pixels for a single trial; vectors initiate at the origin to illustrate the distribution of direction and magnitude. **l,** Flow trajectory of fluorescence in response to reward as mice gained experience with the task. *Top,* Naive mice had irregular responses in the first two days of reward exposure, and at *bottom*, the same mice exhibit smooth waves after 3 weeks of learning reward contingency. See supplementary video 5 for responses plotted.

As in spontaneous conditions reported above, dopamine waves were ubiquitous during task-performance. Notably, reward delivery immediately resynchronized irregular patterns into smooth waves (**Fig. 3g,h**) that had opponent directions depending on task conditions. Specifically, completion of a trial in the instrumental task triggered ML waves (**Fig. 3i,k bottom, Supplementary video 3)**, whereas rewards in the pavlovian task promoted LM waves (**Fig. 3j,k top, Supplementary video 4,** p<0.001 Watson-Wilson test for equality of mean directions in two tasks, n=6 mice for instrumental task, n=8 mice for pavlovian task).

These patterns evolved dynamically with learning: Reward-waves were initially irregular in naive animals but became progressively smooth and directional with experience in task (**Fig. 3l, Supplementary video 5**). The dynamic sculpting of the spatiotemporal patterns by training and task demands ruled out explanations related to the intrinsic anatomy or biophysics of dopamine axons that would constrain the array of observed activation patterns. Thus, we conclude that dopamine waves carry behaviorally relevant decision signals and set out to formalize their precise contribution. In particular, the continuous propagation of dopamine across the striatum both in space and time motivated a revision of standard “temporal-difference” models wherein a single reward-value influences learning about earlier events that are predictive of rewards. We reasoned that these views could be expanded to include “spatiotemporal differences” in which waves carry additional, graded information about structural sub-circuits that are most likely to be responsible for rewards.

## Dopamine waves implement spatio-temporal credit assignment

Our functional interpretation of dopamine dynamics is that the opponent wave trajectories at reward are relevant for spatiotemporal credit assignment. The key inference the animal must make is whether it is in control of the reward-predictive tone transitions, and moreover, which specific contingency applies in the current trial (i.e., distance to run to advance tones). Thus, for mice to preferentially run in the instrumental task (and persist running for long-distance trials), the extent of instrumental controllability should guide reward-evoked dopamine to favor the DMS (i.e. strengthen action-outcome learning). Trial by-trial controllability is partly ambiguous in the task because contingencies were stochastic (drawn from uniform distributions), and mice natively run to varying levels. Nonetheless, we reasoned that task contingencies could still be inferred within trials based on the extent that tone transitions are congruent with locomotion, and dopamine signals can be informed by such congruency.

To formalize this notion, we constructed a multi-agent mixture of experts (MoE) model, extending earlier hierarchically nested corticostriatal circuit models of learning and decision making^(Doya et al., 2002; Frank and Badre, 2011)^ (**Fig. 4a, Supplementary Fig. 7**, see *Methods* for details). At the highest layer (*level 1*) is an expert, putatively corresponding to DMS, that computes the online evidence for action-outcome contingencies and thus task controllability (**Fig. 4a**). Sub-experts within that area (*level 2*) represent specific contingencies (e.g., distance needed to run is short, medium or long) based on previous exposure to the tone transition distributions, learned as a semi-markov decision process via temporal difference learning^(Daw et al., 2006)^. Sub-expert prediction errors (PEs; level 3) occur at tone transitions and are used to compute evidence for (or against) the accuracy of each sub-expert’s prediction. This formulation allows an agent to flexibly adapt behavior based on task contingencies (**Supplementary Fig. 7**)^(Doya et al., 2002; Frank and Badre, 2011)^ and expands the RL account of dopamine to allow both RPE and value signals to be informed by their inferred causal contributions^(Chang et al., 2004; Gershman et al., 2015; O’Reilly and Frank, 2006; Russell and Zimdars, 2003)^.

**Figure 4:**
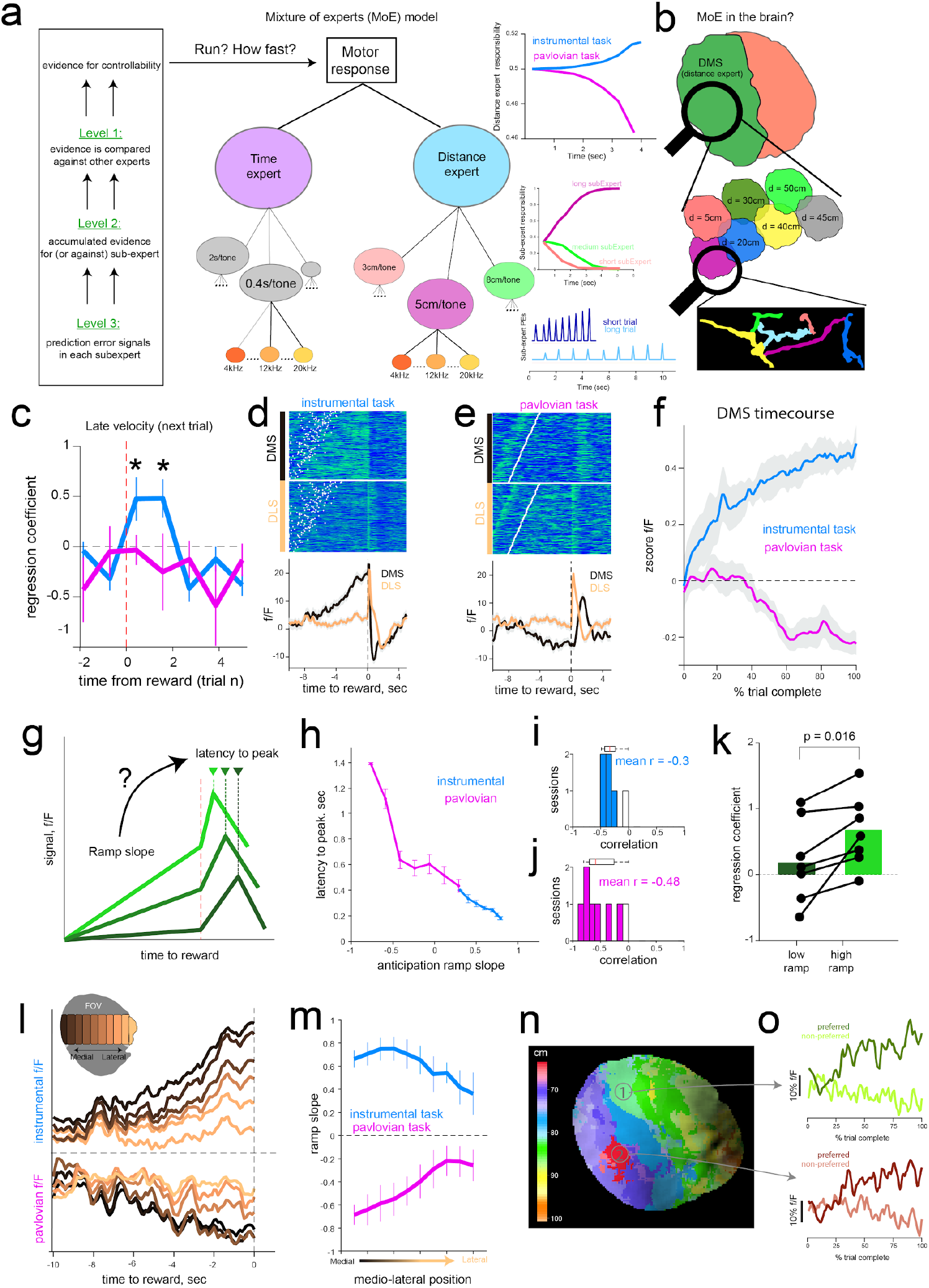
Anticipatory epoch dopamine dynamics reflect inferred controllability and modulate impact of reward on future running in line with a mixture of striatal experts. **a**, Schematic of hierarchical mixture of experts model as a framework for interpreting functional relevance of dopamine dynamics. At the lowest level (level-3), individual states (representing auditory tones) induce reward prediction errors if they misalign with learned contingencies for each “sub-expert”. Prediction errors are accumulated within trials leading to ramps in sub-experts that best describe the current task-contingencies (e.g., short, medium or long distances). At the highest level (level-1), instrumental or Pavlovian “experts” compute the overall (weighted across sub-experts) evidence for instrumental task requirements. These distance-expert responsibility weights accumulate within and across trials to infer controllability of the value function, and are used to adjust model velocities (see Supplementary Fig. 7). **b**, Proposed reflection of MoE signals in striatal dopamine activity, whereby DMS represents overall evidence for controllability (level-1), and is enriched with subregions that represent different instrumental contingencies (level-2). Individual dopamine axon segments signal PEs (level-3) used to accumulate evidence for control. **c**, Multiple regression predicting future running speed of mice in the late phase of the next trial as a function of trial-by-trial wave velocity (in 1 second bins) surrounding the reward from the previous trial. Larger reward-induced waves were predictive of faster mouse running speeds in subsequent trials within instrumental (blue) but not Pavlovian (pink) sessions. Regression coefficients significantly different from zero (blue, asterix p=0.005, two-tailed t-test). Error bars are S.E.M. **d**, Anticipatory and reward response in the medial and lateral DS in a representative instrumental session. White points indicate start of trial. **e,** same format as **d** for pavlovian session. **f,** Aggregate ramping profile during anticipatory epoch for the DMS. Mean activity for each session was z-scored and averaged across mice. Activity in DMS but not DLS showed task-dependent ramping profiles in line with inferred controllability in MoE level 1. Shaded regions represent S.E.M. **g,** Schematic for testing whether anticipatory epochs ramp slope is related to the latency to peak dopamine (peak-normalized within 2sec window following reward) in the outcome epoch. **h**, Results of the relationship from two representative sessions, each from instrumental and pavlovian condition. For both tasks, ramp slope was inversely related to subsequent latency to peak reward response. **i, j** summarize the distribution of correlation coefficients across sessions. **k**, Multiple regression (as in **c**) reveals that impact of reward waves on subsequent trial running speed is stronger when previous trial exhibits large vs small DMS ramps (by median split), in line with credit assignment. **I,** Anticipatory epoch ramps in sample instrumental (*top*) and pavlovian (*bottom*) sessions broken down by ROIs along the medio-lateral axis (inset). **m,** Quantification of ramp slope across sessions. Error bars represent S.E.M. **n,** Local subregions within the dorsal striatum respond preferentially to distinct distance contingencies, reminiscent of sub-expert dynamics (level 2). Contingency specialization map in an example session; color indicates mean distance associated with steepest ramp slopes for each pixel. **o,** Time course of anticipatory ramps in two example subregions for their respective preferred (high ramp trials), and non-preferred (low ramp trials). See Supplementary Fig. 7c for similar pattern of activation in model simulations.

This architecture makes novel predictions at multiple levels which can potentially tie together the separable roles of dopamine during reward pursuit (performance) and learning. First, when reward waves initiate in the DMS (i.e. ML waves in the instrumental task), that region will receive the most credit, and hence mice will be faster to initiate running on the next trial and will persist in doing so until rewards are obtained. Second, the reward wave dynamics should be informed by a trace of which circuit (“expert”) was most responsible for the reward (i.e., which circuit’s predictions were most valuable). We posited that DA dynamics during the tone transitions (anticipatory epoch) could provide such a responsibility signal; that is, the sub-circuit that best predicts the action-outcome contingencies will exhibit increases in dopamine. These levels of dopamine can facilitate mice’s motoric output to be guided most-strongly by that subexpert, while also signaling the degree to which it is responsible for future rewards. We thus hypothesized that DA dynamics during anticipation would impact how reward-waves circulate among striatal subregions and the behavioral expression of running in future trials. Finally, at the most fine-grained level, our model predicts that RPEs should occur at tone transitions to inform the extent of instrumental controllability. In the remaining sections we unpack and test each of these predictions.

The first prediction is that dopamine waves experienced at reward outcome reflects a measure of credit assignment across the striatal experts. ML waves deliver dopamine first to medial subregions (**Fig. 2i,j, 3i,j**), and these DMS-biased signals would selectively strengthen corticostriatal representations for action-outcome contingencies that compete for instrumental control in future trials. As such, we predicted that stronger ML waves at reward would enhance instrumental learning that will drive future running. Indeed, we found a significant correlation between the trial-by-trial magnitude of reward wave and latency to start running on the next trial (n=6 mice, mean r = −0.32, p = 0.0019 two-sided t-test on correlation coefficients). Furthermore, these wave magnitudes predicted the velocity even late in the next trial, 10.2 ±1.4 seconds after the reward response (**Fig. 4c**). The influence of these waves in future-trial behavioral adjustments indicated that they are used for learning functions. Further, these effects were selective to instrumental sessions, indicating that DMS sourced ML waves promote learning about instrumental contingency that is employed for future reward pursuit.

## Anticipatory dopamine ramps provide eligibility for credit assignment

If reward-waves reflect credit assignment, what determines which subregion should receive the credit? Canonical accounts in RL invoke dopamine RPEs that have graded effects on learning depending on “eligibility” signaled by recent MSN activity^(Shindou et al., 2019; Yagishita et al., 2014)^. As noted above, we considered the possibility that local dopamine dynamics during the anticipation epoch themselves signal a coarser measure of eligibility in terms of which subregion was responsible. This trace would be in proportion to the value of the underlying subregion’s predictions, providing a tag for a subregion’s credit.

We thus focused on the activity of dopamine axons during the anticipatory period as mice drew closer to reward. In the instrumental task, we observed a buildup of activity in the DMS (**Fig. 4d,f**), ramping in proportion to the progress to reward^(Hamid et al., 2016; Howe et al., 2013)^. Strikingly, the opposite profile was observed in the pavlovian condition (**Fig. 4e,f**), with decreasing ramps even as the mouse continued to increase licking in anticipation of rewards. These findings are not explained by extant models of DA ramps in accumbens or midbrain, where they have been linked to value functions or RPEs^(Gershman, 2014; Hamid et al., 2016; Lloyd and Dayan, 2015; Morita and Kato, 2014)^, none of which predict opposite profiles across the two tasks.

Instead, we posited that anticipatory dopamine dynamics in the DMS reflects the evidence of agency or controllability, and that subregions within might differentially represent distinct controllable transition functions (which vary from trial to trial). Escalating tones in our tasks provide information about online action-outcome contingency. For example, if tone transitions consistently follow locomotion (as in **Fig. 3d**), they signal evidence for control. The opposite inference can be made in the pavlovian task when tone transitions diverge from locomotion. Respectively increasing or decreasing ramps in the instrumental and pavlovian tasks accumulate in MoE ‘distance’ expert-weights as controllability is confirmed (or contradicted) with each tone transition (**Fig. 4a, right, Supplementary Fig. 7**).

Thus, according to our model, DA ramps do not reflect a monolithic value function, but rather the value of the underlying sub-region’s agentic predictions for reward pursuit, as a marker of that region’s responsibility. Consequently, we argue here that the computational function of dopamine waves at reward-outcome is to assign spatio-temporal credit by delivering dopamine to striatal subregions with different latencies as a function of their graded responsibility signals. This proposal is also motivated by theory and observations that dopamine-mediated plasticity at striatal synapses is strongly attenuated with delayed dopamine release^(Shindou et al., 2019; Yagishita et al., 2014)^.

This credit assignment interpretation makes additional testable predictions at both physiological and behavioral levels. If dopamine ramps during reward anticipation hold persistent information about a sub-region’s prediction accuracy, they should modify the impact of dopamine bursts at reward to focus preferentially on the sub-region with the highest accuracy. As such, striatal areas that ramp with the steepest slopes during anticipation (highest eligibility) should receive a reward response soonest (largest credit, **Fig. 4g**). Indeed, anticipatory ramp slopes across pixels were significantly correlated with the fastest latency to peak fluorescence following reward for both tasks (**Fig. 4h,i**).

Second, if DMS ramps signal responsibility for learning about instrumental control, then trial-by-trial DMS ramp slopes should also modulate the impact of reward waves on next-trial velocity. Indeed, we found that the impact of ML waves on future velocity in the instrumental task (**Fig. 4c**) were dependent on the level of DMS ramps in the previous trial. When DMS ramps were steep, reward waves strongly predicted speeded velocity in subsequent trials; this effect was absent when ramps were weak (**Fig. 4k**, p = 0.016 Wilcoxon signed-rank test, n=6 mice). Together these results suggest that anticipatory dopamine ramps provide a tag for how midbrain driven reward-credit circulate across the striatum to deliver a reinforcement signal for future performance.

Thus far, we have focused on the coarsest division of labor related to the highest level in our model (controllability, level 1), but the agent’s ability to infer control depends on underlying sub-experts that learn distinct action-outcome contingencies (level 2, **Fig. 4a,b**). Such a hierarchical scheme implies that striatal subregions should also differentially ramp for different distance contingencies (**Fig. 4a**). Overall, we observed that DS dopamine ramps are expressed in a gradient across both tasks, with the strongest ramps in the most medial portions (**Fig. 4l,m**). These results are in line with previous work on progressive instrumental specialization of DS on the mediolateral axis^(Thorn et al., 2010)^. Moreover, contiguous territories within the DS exhibit varying ramp profiles for different distance conditions (**Fig. 4n, Supplementary Fig. 8**), with each area expressing the steepest dopamine ramps in preferred set of trials with related distance requirements (**Fig. 4o**). On a trial-by-trial basis, we further observed a significant rank correlation between each pixel’s ramp slope and latency to peak response during reward (mean r = −0.13, spearman’s correlation p<0.001 for all instrumental sessions). These results indicate that the heterogeneously expressed anticipatory ramp gradients across the striatum modulate the spread of reward waves, further strengthening the relationship between eligibility and credit assignment. These findings further support our interpretation that is motivated by MoE account by demonstrating that DMS consists of smaller sub-regions that learn, and express predictions for a variety of potential instrumental contingencies.

These findings led us to ask whether waves organized the response of dopamine axons on the micron scale, and functionally, how evidence for instrumental controllability accumulated in single axon segments. The ramp-like responses we observed at the coarser scale using widefield, one-photon imaging may emerge from trial-by-trial ramps within individual axons, or from a weighted distribution of sharply tuned activation patterns. To directly address these questions, we used 2-photon imaging in two mice to examine the behavior of individual axons in the DMS (**Fig. 5a**).

**Figure 5:**
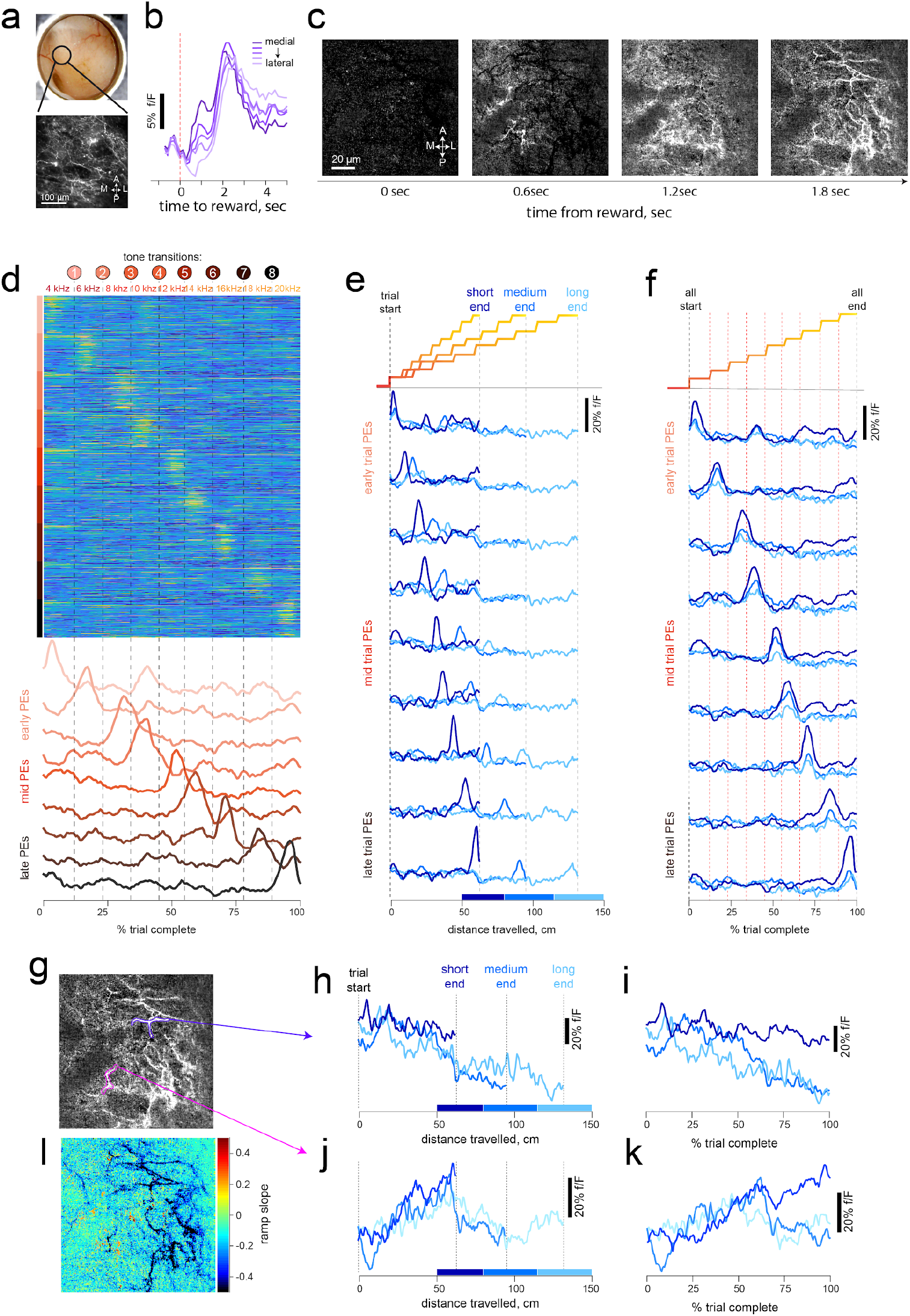
Single dopamine axons show wave-like reward dynamics, tone-specific transients, and distance-dependent ramping during instrumental anticipation. **a**, Schematic of imaged region (*top*), and example field of view of dopamine axons in DMS. **b**, Sequence of frames showing how individual dopamine axons respond to reward. Time relative to solenoid click is shown below each frame. Note the activation of leftmost fibers first, then progressive activation of more lateral axons. **c**, Average time course of reward response from rectangular ROIs equally distributed along the ML axis. **d**, Activity of dopamine segments that respond to tone transitions during anticipation. The activity in each trial is shown as percent trial complete instead of alignment in time. *Top,* heatmap shows the trial-by-trial responses (106 total trials) of groups of pixels in the 2-photon field of view that respond transiently at specific fraction of trial completed. The responses of nine different types are concatenated. *Bottom,* average delta f/F for each type. **E**, Transient response-peaks are not tuned to time or distance run within trial. Each time series is aligned to trial-start and binned into 3 distances: short (50-80cm, dark blue), medium (80-120cm), and long (120-150cm, light blue). Note that the peak location of response arrives at different distances in each trial. By contrast, when aligned to % trial complete, in **f**, the peak response arrives with fixed delay from tone transitions (illustrated at the top for both panels) for all distance contingencies. The transient responses for each tone had larger amplitudes for shorter trials (ie when rewards are predicted to occur sooner, in line with state-dependent reward prediction errors within the lowest level of MoE). **g**, Individual axon segments highlighted to demonstrate example ramp-like trajectories. **h**, Some axon segments ramped downward when aligned to distance travelled or fraction of trial completed as in **i**, as expected at the sub-expert (more localized) level (see Fig 4a and Supp Fig 7). **J** An axon segment that progressively ramps upward only in short distance trials (see Supp Fig 9 for other examples). **k**, same alignment as **i**. **l,** pixelwise map of ramp slope during anticipation.

Similar to our observations at the macro scale, reward delivery recruited dopamine axons in a spatial sequence that was directional (**Fig. 5b,c**), demonstrating that wave-like activation patterns also organize individual axon lattices on the micron scale.

The activity in individual dopamine axons were also modulated during the anticipation epoch. Strikingly, segments of axons transiently responded to auditory tone transition, tiling the full sequence of escalating tones (**Fig. 5d**). The timing of these responses was not affected by distance travelled (**Fig. 5e**), but reliably responded to changes in tone frequency across a variety of distance contingencies (**Fig. 5f**). The systematic tuning of these axons to tone-transitions are consistent with PEs at the lowest level of our model (**Fig. 4a, Supplementary Fig. 7**) that are used to update the online evidence of predictions within each sub-expert. Each tone is represented as a unique state within a sub-expert’s semi-markov process, and PEs arise at tone transitions when they misalign with the predicted distance (or dwell time) until state change. Furthermore, the model predicts larger PEs for state transitions indicative of rewards that will arrive when the distance to run is shorter, due to temporal discounting (**Supplementary Fig. 7**). Supporting this prediction, we observed that tone responses were largest in trials that had shorter distance contingencies, and progressively decreased in amplitude for longer trials (**Fig.5e,f,** mean r = −0.33, and −0.14 n=2 mice; see **Supplementary Fig. 9**). These PE-like responses were distributed throughout the 2-photon field of view, with equivalent fractions of pixels selectively tuned to each tone transition (**Supplementary Fig. 9**).

We also noted that contiguous segments of axon lattices had single trial ramps that were either upward (**Fig. 5h,i**) or downward (**Fig. 5j,k**) as mice get closer to reward. Instead of tuning to tone-transitions reported above, these dopamine ramps were selectively expressed for different contingencies, ramping to varying extents depending on the required distance in separate trials. Together these results provide evidence for two, simultaneous classes of nearby dopamine axon segments (**Supplementary Fig. 9**) used for sub-expert computations: transient PE signals that respond to state transitions, and ramping segments that accumulate evidence for controllability as predicted by a sub-expert.

## Discussion

Our report of dopamine waves provides the earliest evidence for a foundational organizational principle of dopamine axons that correlate activity within functionally related striatal boundaries. In the cortex, travelling waves have been described to facilitate (or constrain) computations that are topographically organized^(Bringuier et al., 1999; Muller et al., 2014; Nauhaus et al., 2009)^. Similarly, we interpret the computational significance of dopamine waves as orchestrating dopamine release to striatal subregions that exhibit a graded functional specialization on the medio-lateral axis^(Barbera et al., 2016; Klaus et al., 2017; Thorn et al., 2010)^. Thus, waves are a natural candidate for solving the spatiotemporal credit assignment problem when multiple, topographically organized striatal actors/sub-experts compete to guide action selection across multiple levels of abstraction^(Collins and Frank, 2013; Doya et al., 2002; Frank and Badre, 2011)^. We used a very simple task to manipulate reward and sensory statistics, requiring mice to resolve ambiguity about instrumental contingency by comparing predicted and actual tone transitions. Consistent with the MoE account, wave directions during reward were sensitive to controllability of task structure, and -- only in the controllable task -- dopamine waves were related to future behavioral adjustment on a trial-by-trial basis.

We also describe anticipatory epoch ramping dynamics that appear to signal the value of a subregion’s prediction about reward contingency. These dynamics may serve a dual purpose. First, they could promote online behavioral flexibility (e.g., optimize reward-rate and minimize energetic costs) according to the predictions of the most accurate subregions during reward pursuit. Second, these activity patterns would also signal which subregion was most responsible for behavioral output and hence provide a low dimensional tag for responsibility (akin to an eligibility trace in RL^(Singh and Sutton, 1996)^), which would then allow for reward-driven RPEs to preferentially credit the appropriate subregion and the eligible MSNs within it. While the two functions are not mutually exclusive, our data provide strong support for the second interpretation: On a trial-by-trial basis, the degree of ramping across regions was related to the latency to reward peak elicited by the wave, and the combination of ramp slope and wave magnitude was predictive of subsequent-trial behavioral adjustments. These findings accord with views that dopamine signals can have different functions during reward pursuit and outcome, which can be gated by local microcircuit elements that regulate plasticity windows^(Berke, 2018; Bradfield et al., 2013; Franklin and Frank, 2015; Morris et al., 2004; Threlfell and Cragg, 2011)^. Moreover, we also interpret transient and localized RPEs during reward pursuit as facilitating inference about the current task state (i.e., determining credit), whereas RPEs during reward itself facilitates reinforcement learning; a dual operation that can also be gated^(Franklin and Frank, 2015; Gershman et al., 2015; Redish et al., 2007; Schoenbaum et al., 2013)^. Put together, the synthesis of our data and computational simulations imply that dopamine signals are spatio-temporally vectorized during both epochs, tailored to underlying region’s computational specialty.

Although DMS dopamine support the computations of the ‘Distance’ expert in the MoE, one limitation of our study is that we did not identify or assess the dopamine dynamics with properties of the ‘Time’ expert. Many studies investigating RPEs involve classical conditioning in which temporal representations are evident in the midbrain^(Hollerman and Schultz, 1998; Pan et al., 2005; Soares et al., 2016)^, and ramping signals related to timing may be present in other regions upstream of the DA system^(Brown et al., 1999; Hazy et al., 2010; Mello et al., 2015)^. Nonetheless, even without timing experts, the DMS expert in our model could behave similarly across the two tasks if it simply evaluates evidence for agency relative to some prior expectation about control.

Another limitation of our study is that we did not deduce the origin of dopamine waves, which may be inherited from sequential firing of midbrain dopamine cells that have a topographical projection pattern^(Lerner et al., 2015)^. To date, such dynamics have not been reported in the literature, potentially because limited studies investigated the activity of a large population dopamine neurons simultaneously^(Engelhard et al., 2019)^. Another likely mechanism may involve local sculpting of dopamine release within the striatum. Wave-like patterns have been reported in neocortex^(Kasanetz et al., 2008; Mohajerani et al., 2013)^ and striatal cholinergic interneurons^(Rehani et al., 2019)^, both of which can potently regulate dopamine axon activity^(Cachope et al., 2012; Krebs et al., 1991; Threlfell et al., 2012)^. Moreover, dopamine waves at reward outcome may also be a consequence of the interaction between primed excitability of dopamine axons during the anticipatory epoch and midbrain-sourced synchronous reward bursts. The combination of these two patterns may produce sequential activation that propagates across the striatum in proportion to expressed ramp during anticipation.

## Methods

### Animals and Surgery

All procedures were conducted in accordance with the guidelines of the NIH and approved by Brown University Institutional Animal Care and Use Committee. We used 17 DAT-cre mice (9 females, 8 males; *Jax Labs # 020080*) that were maintained on reversed 12hr cycle and all behavioral training and testing was performed during the dark phase. To achieve selective expression of GCaMP6f in dopamine cells, we followed standard surgical procedures for stereotaxic injection of cre-dependent virus. Briefly, mice were anesthetized with isoflurane (2% induction and maintained at 0.75-1.25% in 1 liter/min oxygen). To attain widespread infection of dopamine cells throughout the midbrain, we drilled two burr holes above the midbrain (−3.2mm AP, 0.4mm and 1.0mm ML relative to bregma) and injected 0.1-0.2µL of AAV-syn-Flex-GCaMP6f at two depths per burr hole (3.8 and 4.2 mm relative to brain surface). We next secured a metal head post to the skull and implanted an imaging cannula over the ipsilateral dorsal striatum. The cannula is a custom fabricated stainless-steel cylinder (Microgroup; 3mm diameter and 2.5-3mm height) with a 3mm coverslip (CS-3R, Warner Instruments) glued at the bottom with optical adhesive (Norad Optical #71). To insert the cannula into the brain, a 3mm diameter craniotomy was first drilled over the striatum (at bregma, centered on 2.0mm ML), and then gently removed the dura and slowly aspirated the overlying cortex until white colossal fibers were clearly visible (∼0.8-1.2mm from brain surface). These fibers were also gently aspirated layer by layer until the underlying dorsal striatal tissue was uniformly exposed. A sterile imaging cannula was progressively lowered until the coverslip contacted striatal tissue uniformly. Dental cement was applied to secure implant to the skull and mice received a single dose of slow-release buprenorphine and allowed to recover for 1-2 weeks with post-operative care.

### Behavioral Training

After full recovery from surgery, mice underwent 2-3 days of habituation in operant chambers outfitted with a 3D printed wheel (15 cm diameter), audio speakers and a solenoid-gated liquid reward dispenser. Following acclamation, mice were water-restricted, receiving 1ml/day in addition to water earned during task performance. We used custom LabVIEW scripts to control operant boxes during training and testing in behavioral tasks. In the first stage of training, mice received non-contingent rewards that were delivered randomly (3-15 second apart, uniform distribution) for 3 consecutive days. Next, training in a “pavlovian” task began, wherein rewards were delivered after a variable delay from trial start. The start of each trial is signaled by the onset of a 4.3kHz tone that continued to escalate in frequency in proportion to fraction of trial completed. We used nine different frequencies that were selected to minimize harmonic overlap; 4.3kHz, 6.2kHz, 8.3kHz, 10kHz, 12.4kHz, 14.1kHz, 16kHz, 8.4kHz, 20kHz. Across trials, the duration to wait for reward is randomly drawn from a uniform distribution (4-8 seconds). At the end of a trial, the auditory sound is turned off, and the solenoid delivered 3µL of water reward to a spout in front of the mouse. Licking behavior is detected using capacitive touch sensors (AT42QT1010, Sparkfun). In some catch trials, the initial 4.3kHz tone turned off after 0.5s and the mouse did not have continuous information of progress to reward. For clarity, we only focused on escalating-tone trials. The next trial started after a variable inter-trial-interval of 3-8 seconds. After 2-3 weeks of the pavlovian task, activity of dopamine axons in the striatum was imaged in a test chamber with a widefield and 2-photon imaging system. The same animals were then further trained on a distance-variant of the same task, where reward delivery is now contingent on mice running on the wheel. Mice were exposed to the “instrumental” task in training chambers requiring them to run on the wheel to traverse linearized distances, also randomly selected from a uniform distribution (50-150cm). Progress to reward was indicated by the same tone frequencies, and the angular position of the wheel recorded using a miniature rotary encoder (MA3A10250N, US Digital). All behavioral data is digitized and stored to disc at 50Hz.

### Widefield and two-photon imaging

All imaging was performed using a multi-photon microscope with modular laser-scanning and light-microscopy designed by Bruker/Prairie Technologies. For widefield imaging, we used a full-spectrum LED illumination with FITC filter cassette for illumination at 470nm and detection centered at 530nm. Images were acquired using a CoolSnap ES2 CCD camera (global shutter, Photometrics) and synchronized with behavioral events through TTL triggers. All widefield images during behavioral tasks were acquired with a 4X objective (Olympus), 100ms exposure (10Hz) and 8X on-camera binning to achieve a sample resolution of 40µm/pixel (unless indicated otherwise). Two-photon microscopy was performed using a 20X air objective (Olympus) on the same imaging platform with a femtosecond pulsed TiSapphire laser source (MaiTai DeepSee, 980nm power measured at objective was 20-50mW) that was scanned across the sample using a resonant (x-axis) and non-resonant (y-axis) galvanometer scanning mirrors. Returning photons were collected through an imaging path onto milti-alkali PMTs (R3896, Hamamatsu), and recorded frames were online-averaged to achieve a sampling rate of 10-15Hz.

### Data Analysis and statistics

All images were processed with custom routines in MATLAB. Each session is preprocessed for image registration, and alignment to behavioral events based on triggers. Movement artifacts and image drift in the XY plane were corrected using rigid-body registration using a DFT-based method^(Guizar-Sicairos et al., 2008)^. To cluster the activity of dopamine axons, we used the K-means algorithm in MATLAB. To compute robustness of clustering results, we used the adjusted rand-Index measure which computes the similarity of two clusters based on the probability of member overlap (corrected for chance; 0=random clusters, 1=exact same membership). To examine how robust the clustering results were, we re-clustered the same dataset 100 times in K-means using random initialization and varied cluster limits. We compared the extent that pixels were re-clustered into the same group using the adjusted rand-index as an indicator of robustness of underlying structure of the data that produced clusters (see Supplementary Fig. 4). To additionally test the extent spatial relationship between the pixels, or their temporal relationship influenced the identified clusters, we repeated the same analysis but shuffled the spatial or temporal relationships between the pixels. To estimate the optimal number of clusters within each dataset, we computed the Bayesian information criterion (BIC) on the K-means algorithm.

We characterized flow patterns in dopamine waves by adapting standard optical flow algorithms in machine vision that are adapted for imaging of fluorescence signals^(Afrashteh et al., 2017; Mohajerani et al., 2013; Townsend and Gong, 2018)^. Briefly, flow trajectories were computed for any two successive frames as a displacement of intensity across the pixels in time. This method allows us to evaluate a pixel-by-pixel velocity vector fields that summarizes the direction and strength of flow at each pixel. While there are multiple methods to achieve this calculation^(Afrashteh et al., 2017; Townsend and Gong, 2018)^, we adapted a combined Global-Local (CGL) algorithm^(Bruhn et al., 2002; Liu, 2009)^ that combines the Lucas-Kanade and Horn-Schunck methods. The frame-by-frame vector fields calculated using the CGL method was further processed to extract sink and source locations and also flow trajectories across multiple frames (**Fig. 2a, bottom**). The frame-by-frame flow magnitude for each frame (or flow-velocity, with units of mm/second) is computed by averaging the length of vectors at each pixel (e.g. **Fig. 2b, red**). The locations of sinks or sources were estimated based on local vector orientations: i.e sinks are points of inward flow, whereas sources are points of outward flow. We estimated the pixel-wise likelihood of sinks and sources by simply computing the divergence of the vector field in each frame (“*divergence”* function in MATLAB, **Supplementary Video 2**). The flow trajectory across frames were calculated from vector fields using the “*stream3*” function in MATLAB from seeded pixels (e.g. **Fig3l**).

For alignment of fluorescence time series, DMS and DLS masks were defined using one of three methods: i) manual drawing, ii) boundaries using cluster results (as in Fig. 1i, top) and iii) uniformly spaced ROIs on the mediolateral axis (as in Fig. 4l, inset). Each animal performed multiple behavioral sessions, and we used one session per animal (n=6 mice in instrumental task, and n=8 mice in pavlovian task) that had the largest Δf/F deviations to avoid results from being dominated by a few animals.

To determine the influence of reward-wave on behavioral performance on the next trial, we performed a multiple regression predicting the running velocity of mice late (i.e. 75-100% of trial complete) in the next trial based on reward-aligned wave magnitude (1-sec bins, **Fig. 4c**). To determine whether DMS ramp slopes influenced how last-trial wave outcome, on the next trial, we conditioned this analysis on the ramping profile in the DMS, median-split into low and high ramp conditions (**Fig 4k**). We evaluated the correlation between the ramp slope and latency to peak by first peak-normalizing the reward response in 2-sec window and finding the time index (after reward) for which the fluorescence signal reached peak levels. To examine whether anticipatory dopamine ramps had a preference for different distance conditions (**Fig. 4n,** also see **Supplementary Fig. 7**), we sorted the trials based on the expressed ramps in each pixel and averaged the distance contingency in trials with top 90% ramping.

TIFF stacks of 2-photon images of dopamine axon segments were also pre-processed for registration and alignment with behavioral data. To draw ROIs of these segments for assessing organization of responses (**Supplementary Fig. 1**), we followed the Howe and Dombeck^(Howe and Dombeck, 2016)^. Otherwise, we generally used pixel-wise analyses.

### Computational model

We modeled mouse behavior using a mixture of experts / multi-agent RL architecture^(Frank and Badre, 2011)^, extended here to accommodate the sequential tone structure with semi-markov dynamics^(Daw et al., 2006)^. We modeled the two task structures as separate “experts” that learned a value function *V* as a function of either elapsed time as in classical temporal difference learning applied to Pavlovian condition, or as a function of distance travelled. Because mice were trained on both time and distance tasks, multiple sub-experts (representing clusters in mediolateral coordinates of striatum) were pre-trained for 2000 trials to span a range of contingencies (e.g., 400ms, 600ms, or 800ms per tone transition; or 5, 10 or 15cm). For simplicity, we modeled the task with discrete sub-experts that specialized on (had been preferentially exposed to) particular times/distances. However, one can easily generalize the framework to the continuous case (e.g., using basis functions^(Ludvig et al., 2008)^) and the discrete space can be modeled with arbitrary resolution by simply increasing the number of sub-experts. Moreover, various models have shown that prediction errors can be used to segregate learning of different latent task states^(Collins and Frank, 2013; Gershman et al., 2015)^.

#### Subexpert and expert learning

The value function for each time sub-expert *s* estimates the discounted future reward V^s^(*Xi,t*) = r(t) + *γ*V^s^(*X_i,t +1_*) and was trained via temporal Differences^(Sutton and Barto, 1998)^ based on reward prediction errors *δ(X_i,t_) = r(t) + *γ*V(X_i,t +1_) - V(X_i,t_)*. Each auditory tone was modeled as a distinct state *X_i,t_* or X_i,d_ with semi-markov dynamics. That is, the onset of each tone *i* would advance the state vector to the corresponding position even if the tone occurred earlier or later in absolute time/distance. Thus the value function learned for each sub-expert was tied to the current state (tone) and the (discretized) dwell time (*t)* or distance (*d)* since it has been entered, and not to the absolute time or distance that passed from the onset of the first state. This semi-markov process was based on the assumption that the tone stimuli induce a neural state representation upon which TD is computed^(Daw et al., 2006; Ludvig et al., 2008)^ and evidence that rodents are endowed with such a rich state representation^(Gardner et al., 2018)^. The value function was learned by adjusting weights in response to the X state vector, with *V(X_i,t_)= w_t_ X_i,t_* and *w_t_ ← w_t_ + *α* δ(t), α* where is a learning rate. The distance experts were trained analogously, but with the X vector advancing with each (discretized) distance step rather than passive time. Thus if the agent stopped moving, the *X_i,d_* vector remained constant until it moved again, and if it moved faster than usual, the *X_i,d_* vector would advance to later states accordingly. We fixed *α* =0.25 and *γ* =.95 for all experts but verified that the patterns were robust to other settings.

#### Performance and inference

After learning, the on-line evidence (responsibilities, fig 4B and supplemental fig7, modeling the ramps) for each sub-expert was computed as an approximation to the likelihood of the trial-wise tone transitions for that sub-expert. We adopted a hybrid Bayesian-RL formulation^(Frank and Badre, 2011)^. From a Bayesian perspective, the attentional weights for each expert can be evaluated by computing the posterior probability that each expert encompasses the best account of the observed data x: *P(s|x) = P(x|s) P(s) / Σ P(x|s_i_) P(s_i_).* Thus the evidence for each expert is computed by considering its prior evidence *P(s)* and the likelihood that the observed tone transitions or rewards would have been observed under the expert’s model *P(x|s)*, relative to all other experts. For example, if there was a low probability for a tone transition at a particular moment under a given expert, then the likelihood of that observation given the expert’s model is low. Once the posterior evidence for each expert is computed, one can then apply Bayesian model averaging to allocate attentional weights to each expert in proportion to their log evidence.

Rather than a fully Bayesian realization, we instead implemented an RL approximation that may more directly relate to corticostriatal DA mechanisms^(Frank and Badre, 2011)^. Instead of computing the likelihood directly, expert responsibility weights were assigned such that experts with the smallest Bellman errors *δ_s_* accumulated the most weight. In particular, the responsibility weight for each subexpert *ω ‘_s_* was decremented when the corresponding sub-expert experienced a reward prediction error: *ω ‘_s_ ← *ω* ‘_s_ − δ_s_,* where δ_s_ is the positive reward prediction error according to the corresponding sub-expert’s value function given state vector X. (Similar results hold if using |*δ_s_*| instead of only positive RPEs to decrement expert weights). Intuitively, experts with more prediction errors are less likely to have been responsible for the outcome (tone transition or reward). These responsibility weights were then normalized relative to all sub-experts as an approximation to the log evidence for a given subexpert: *ω_si_ =* exp(*βωsi*) */ Σ j*exp(*βωsj*), where *β*is an inverse temperature parameter. Thus, in contrast to standard RL in which RPEs reinforce actions that yield rewards, during inference, more frequent Bellman errors for a given subexpert are indicative that it is less responsible for observations compared to subexperts that have minimal error. Such a scheme is compatible with extant models that use reward prediction errors for state creation and inference separate from reinforcement per se^(Collins and Frank, 2013; Frank and Badre, 2011; Gershman et al., 2015; Redish et al., 2007)^. We posited that these RPEs correspond to the phasic events observed at tone transitions in the 2p imaging data. The accumulation of these responsibility weights were posited to relate to the 1p imaging data in discrete sub-regions of DMS.

Finally, a second-level task selection process was implemented to arbitrate responsibility between the overall distance expert and overall time expert (each of which constituted a weighted combination of their subordinate experts). This inference process was identical to that for the sub-experts, with responsibility updated based on their experienced prediction errors: *ω ‘_D_ ←*ω*‘_D_ − δ_D_*, where *ω‘_D_* is the accumulated responsibility of the distance expert based on its reward prediction errors, *δ_D_= r(t) + γV_D_(t+1) - V_D_(t)*. The value function for the distance and time experts V_D_ and V_T_ are in turn weighted averages according to the inferred responsibilities of the subordinate experts within each structure: *V_D_(t)= Σ *ω*_sD_V_SD_*(t) and *V_T_ = Σ*ω*_sT_V_ST_*(t). Similarly, the value function of the agent as a whole is the weighted average value function across the two experts *V(t) = *ω*_D_ V_D_ (t)+ *ω*_T_ V_T_*(t). These responsibility weights for each task structure were again normalized across tasks, *ω_D_ = e^βw‘D^ / e^β w‘D^+ e^βw‘T^*.

For each distance or time, 100 test trials were run with 10 tones each and an inter trial interval was randomly drawn from 5-15s. The agent as a whole selects actions in terms of speeds to run for a period of time at each tone transition or after it has completed it’s previous running. Speeds were selected in proportion to the inferred responsibility of the DMS expert, together with some stochasticity: speed(t) = 5\**_D_* (t)-0.5)) + *ϵ*, where *ϵ* was drawn from a uniform distribution with a mean of 3. Stochasticity facilitates the agent ability to disambiguate distance from time tasks within a trial (a constant speed would equate the prediction errors for the two tasks given appropriate sub-experts). Increasing speed with inferred DMS expert responsibility **ω*_D_* allows the model to capture the increased running with instrumental task structure (Supplementary Fig 7). More detailed investigation of how speeds may be optimized according to reward/effort/delay tradeoffs will be examined in future work.

## Acknowledgements

We thank Matthew Nassar, Joshua Berke, Peter Dayan, Theresa Desrochers, for valuable discussion of the project and feedback on an earlier version of the manuscript, and members of the Frank and Moore laboratories for feedback at various stages of the project. We also thank Ines Belghiti and Aneri Soni for help with adapting the MoE model implementation to the tone task. GCaMP6 was developed and generously made available by the HHMI Janelia Farm Research Campus GENIE Project. This work was supported by HHMI Hanna Gray Fellowship to AAH, R01MH080066 and NSF grant 1460604 to MJF, and awards from Carney Institute for Brain Sciences and Dean’s office to CIM.

## Contributions

AAH designed and performed the experiments, analyzed the data, and applied computational model and wrote the paper. MJF designed and supervised the study, developed computational model, and contributed to writing and revising of the paper. CIM designed and supervised the study and revised the paper.

## Competing Interests

The authors declare no competing interests.

## Data and code availability

All data and code is available from corresponding author(s) upon reasonable request.

## Materials and Correspondence

Correspondence and material request is to be addressed to Arif Hamid, Christopher Moore or Michael Frank.

---

**Supplementary Fig. 1:**
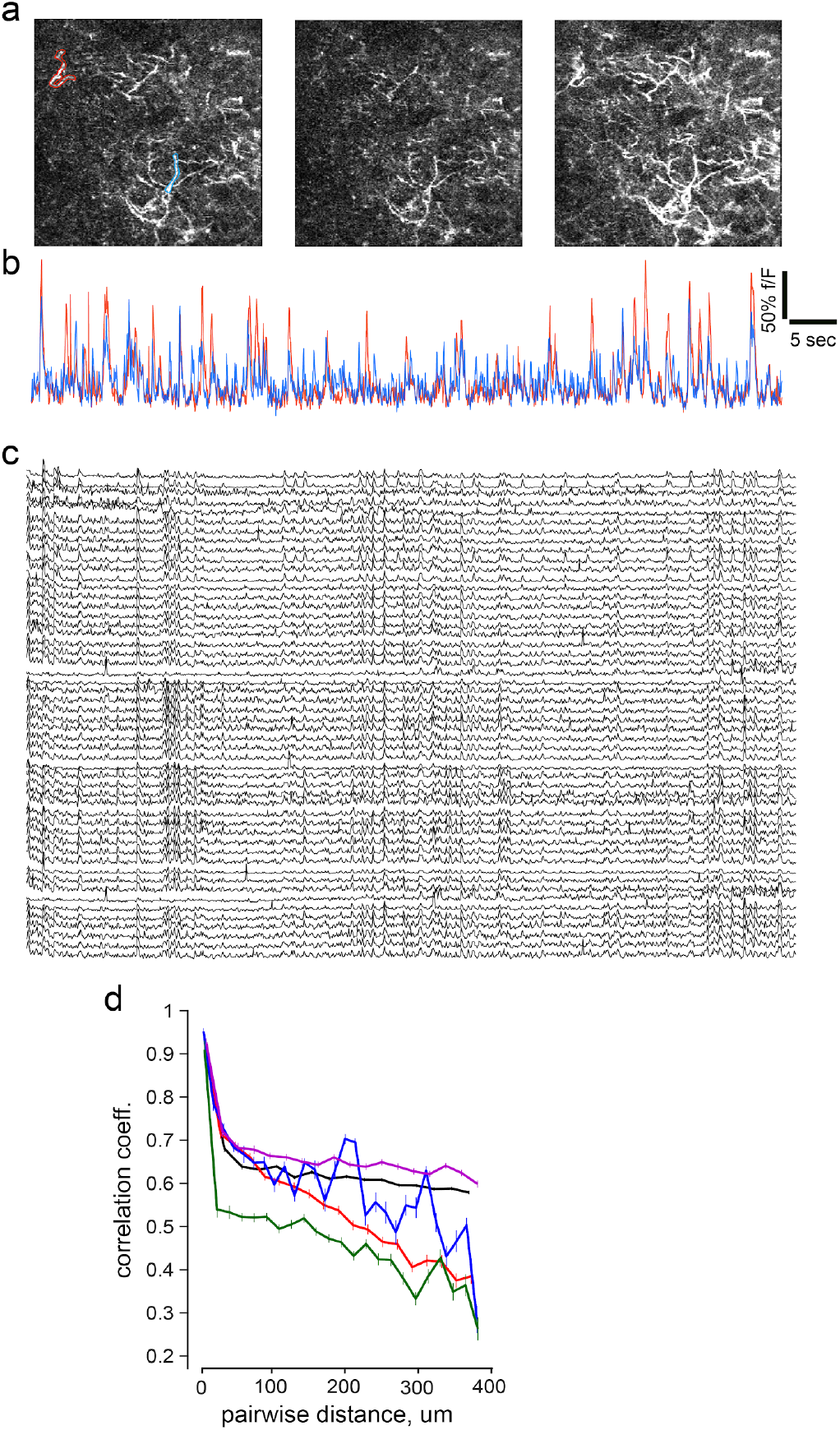
Individual dopamine axons also exhibit decorrelated activity patterns. **a**, Example frames illustrating that different portions of axon laticies are activated asynchronously. **b,** Representative timeseries of fluorescence from two axon segments outlined in blue and red at **a**. **c,** Additional examples of activity in dopamine axon segments. Data is organized such that rearby axons are plotted closer. **d,** Quantification of correlation between the session wide timesereries of axons based anatomical distances. Note that nearby axons are highly correlated, but they exhibit a distance dependent falloff as reported in Fig. 1g, although a different anatomical scale.

**Supplementary Fig. 2:**
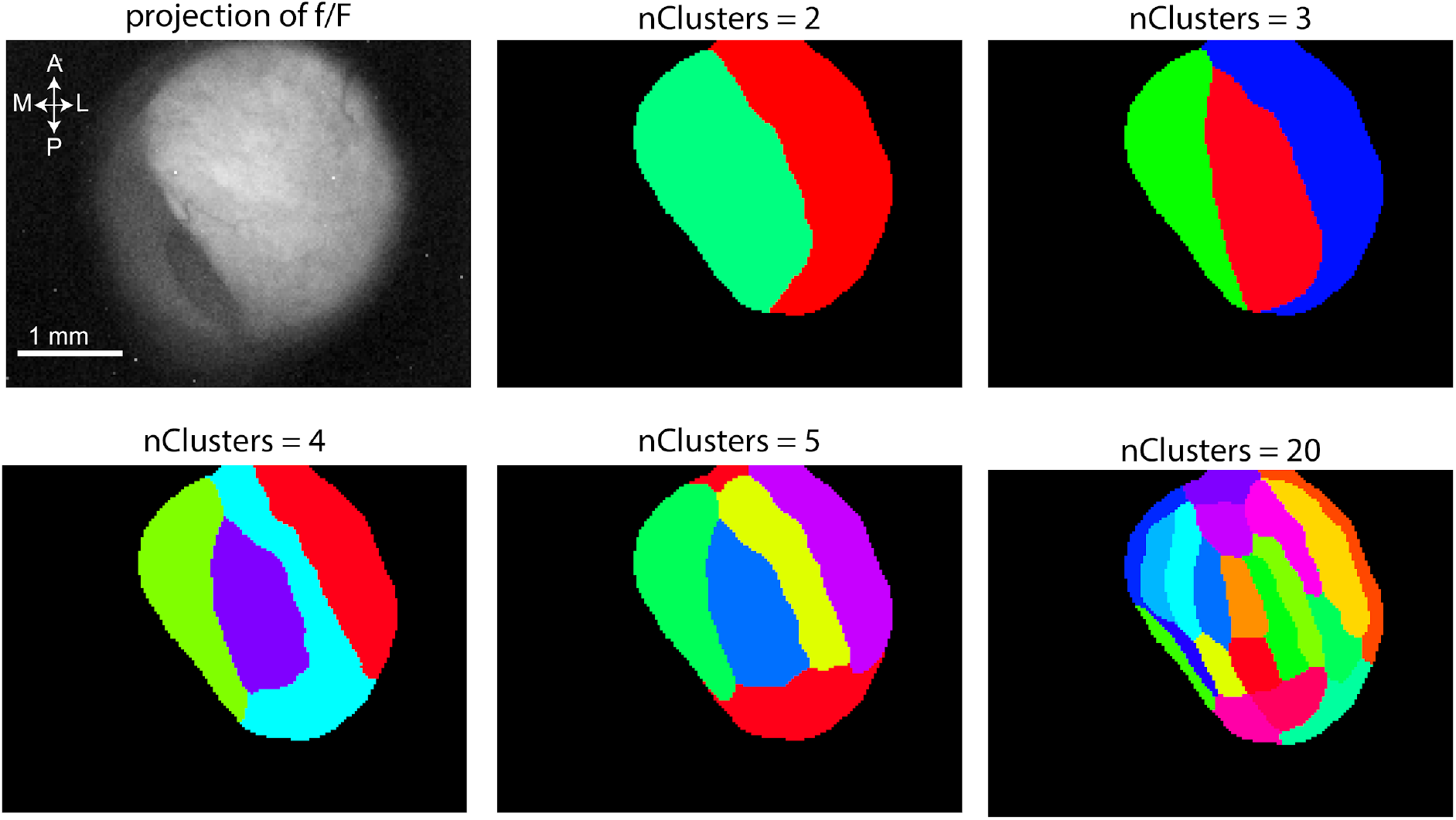
Dopamine axon activity clusters into hierarchical domains. Data from one session, mean projection of fluorescence is displayed at the top left. Progressively increasing cluster limits identifies contiguous striatal subregions that decompose into sub-clusters.

**Supplementary Fig. 3:**
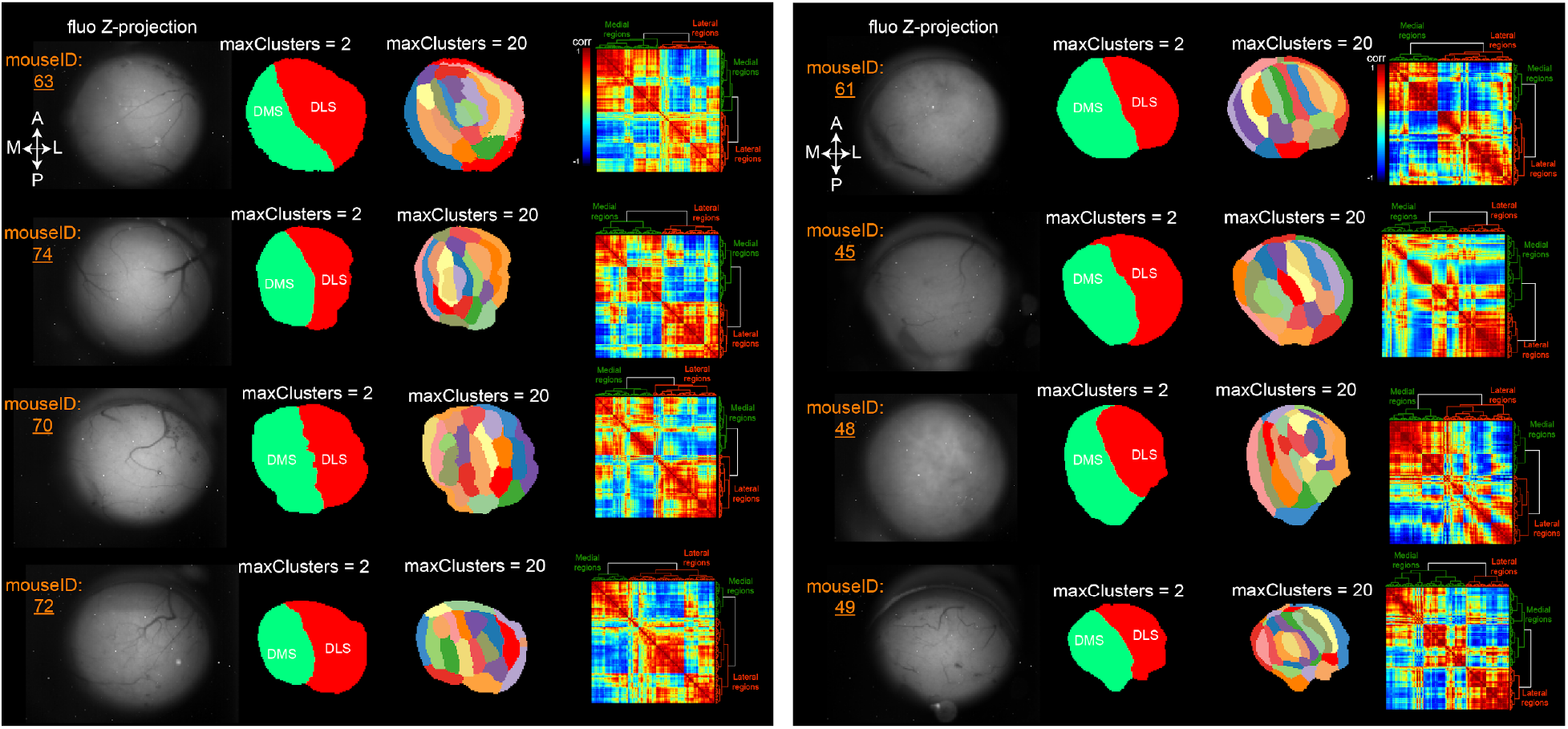
Clustering patterns in all 8 mice. We provide the K-means cluster of each of the animals examined. Plotting format follows panels in Fig 1.

**Supplementary Fig. 4:**
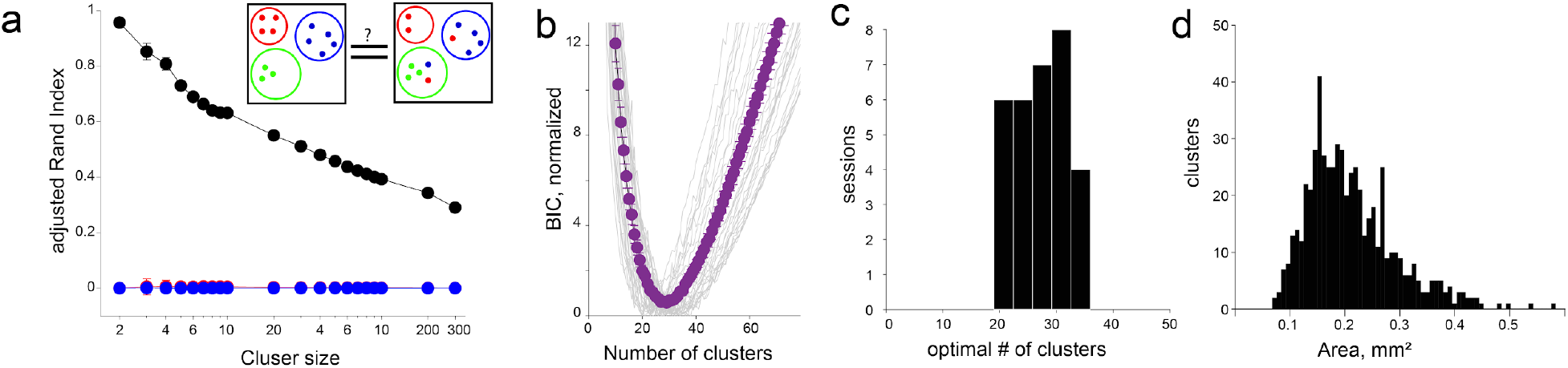
Cluster patterns are robust. **a,** Adjusted rand-index score of cluster patterns determined using the K-means for re-clustering (black) or shuffling the temporal (red) or spatial(blue) indices of pixels. Results are shown for 100 reclustering iteration with or without shuffling. Note that randomizing the temporal or spatial relationships of fluorescence activity results in random clusters. **b,** BIC score for K-means results all sessions examined (n=31 sessions, 8 mice; gray), and average (purple). **c,** Distribution of optimal number of clusters identified using the BIC metric. **d,** Distribution of areas of identified clusters across all mice.

**Supplementary Fig. 5:**
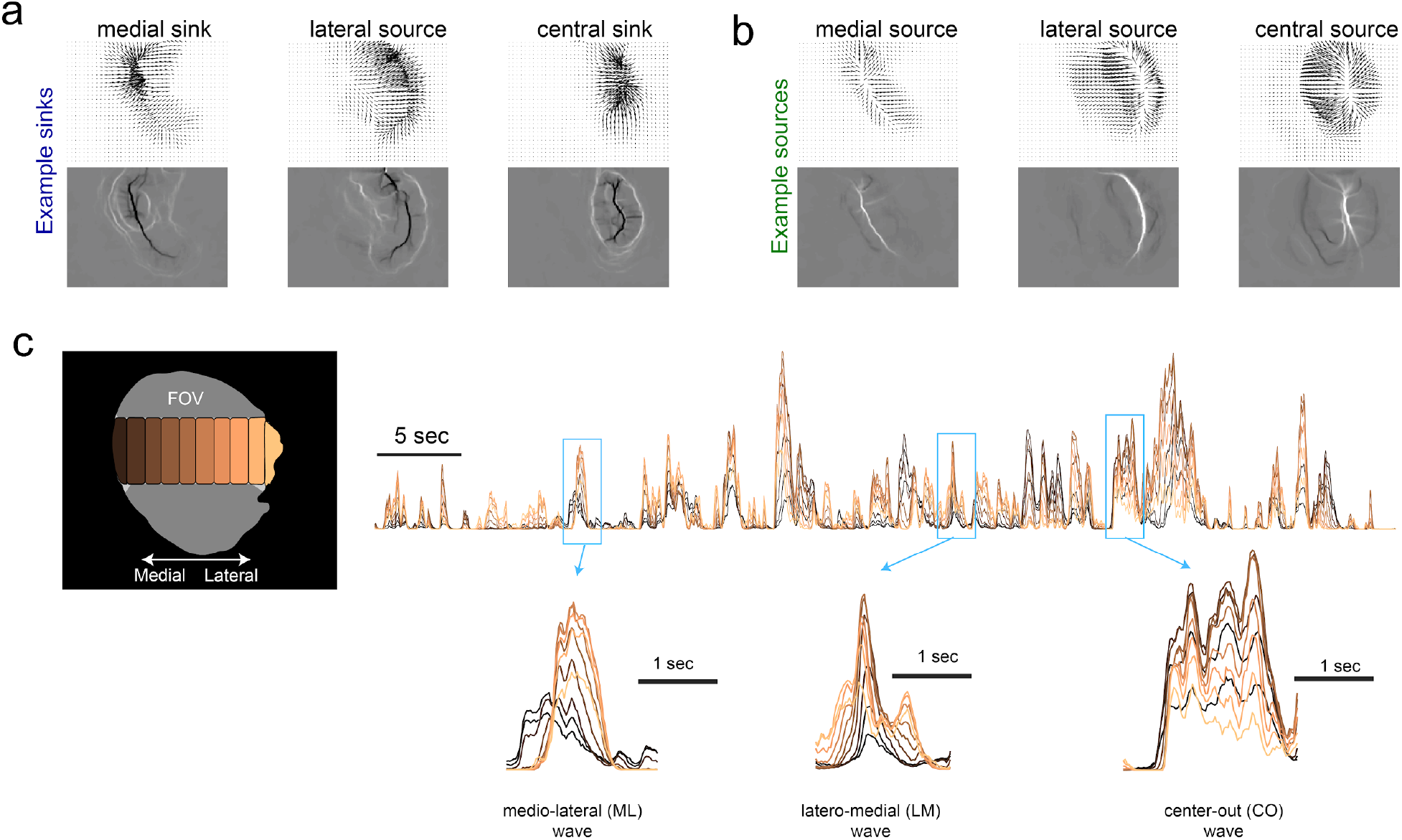
Local sources and sinks initiate and terminate dopamine activity, delivering temporally delayed dopamine to striatal subregions. **a,** Flow pattern (*top*) amd divergence map (*bottom*) for sinks that are clustered in medial, lateral or central striatal regions. b, same format as a, for source locations. c, Time Course of activity across the mediolateral gradient for a one minute recording epoch. Blue boxes focus on transient events that were produced by ML, LM, or CO waves that deliver dopamine to different parts of the dorsal striatum with relative lags.

**Supplementary Fig. 6:**
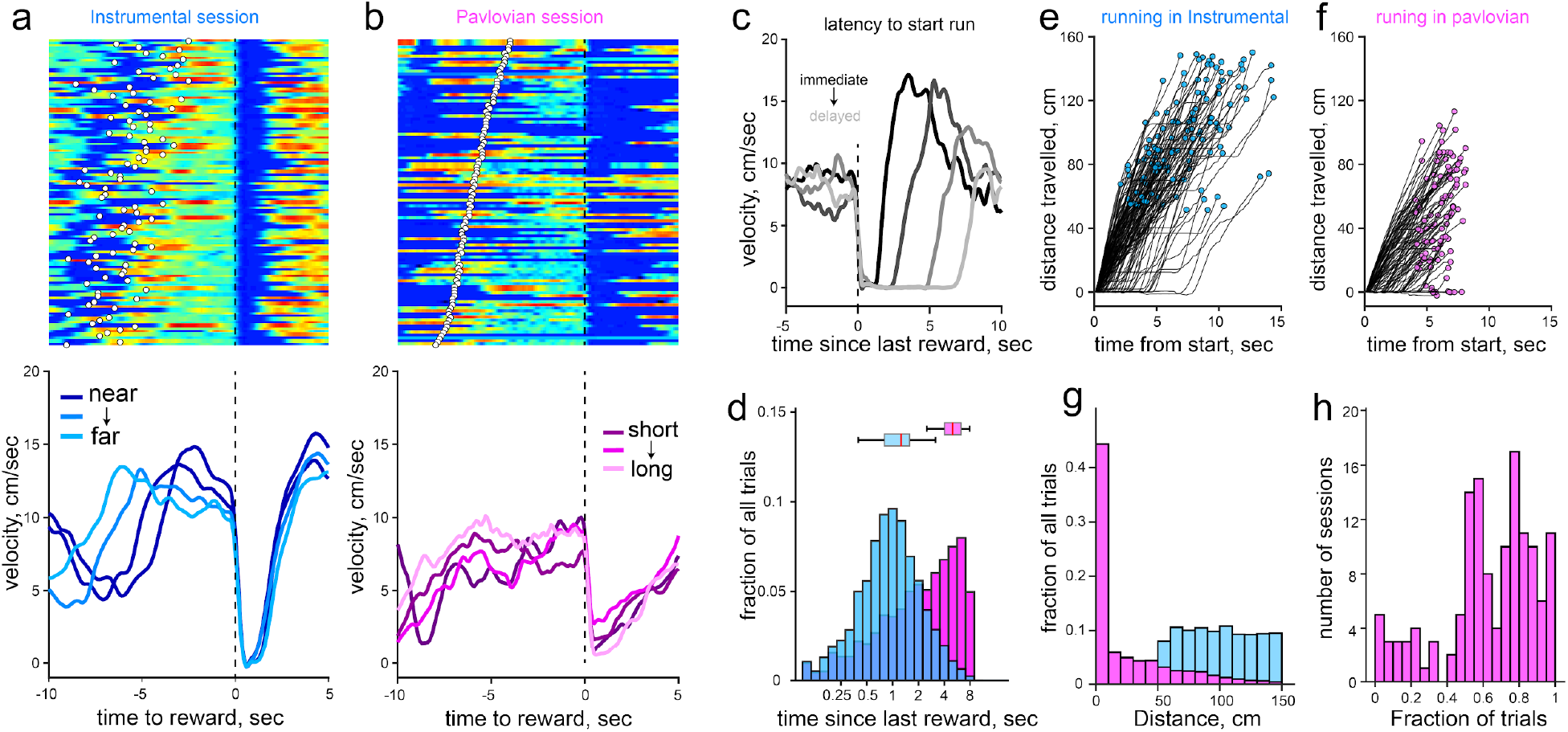
The running behavior of mice is more structured and goal-directed in the instrumental task. **a,** Example velocity profile for an instrumental session. *Top* shows the trial by trial velocity aligned to the end of trial (reward receipt) White dots indicate the start of the trial. *Bottom* illustrates mean velocity trajectory for different distance contingencies. **b,** Same format as **a** but for pavlovian session. Note that the running behavior of the mouse is disorganized relative to task events (quantified in later sessions). **c,** Example session showing variability the latency to start running on the next trial. **d,** We quantified this latency for training sessions across all mice and observed a significantly shorter latency to initiate next trial running. X-axis is displayed in log scale. **e,** Single trial trajectories of position from trial start during instrumental sessions, and pavlovian sessions in **f**. Circles denote mouse position at the end of a trial. **g,** Overall, mice ran less distance that the requirement in instrumental sessions (i.e. 50-150cm), and **h**, mice chose not to run at all in a significant fraction of trials during the pavlovian task (note that running is required for rewards in instrumental sessions).

**Supplementary Fig. 7:**
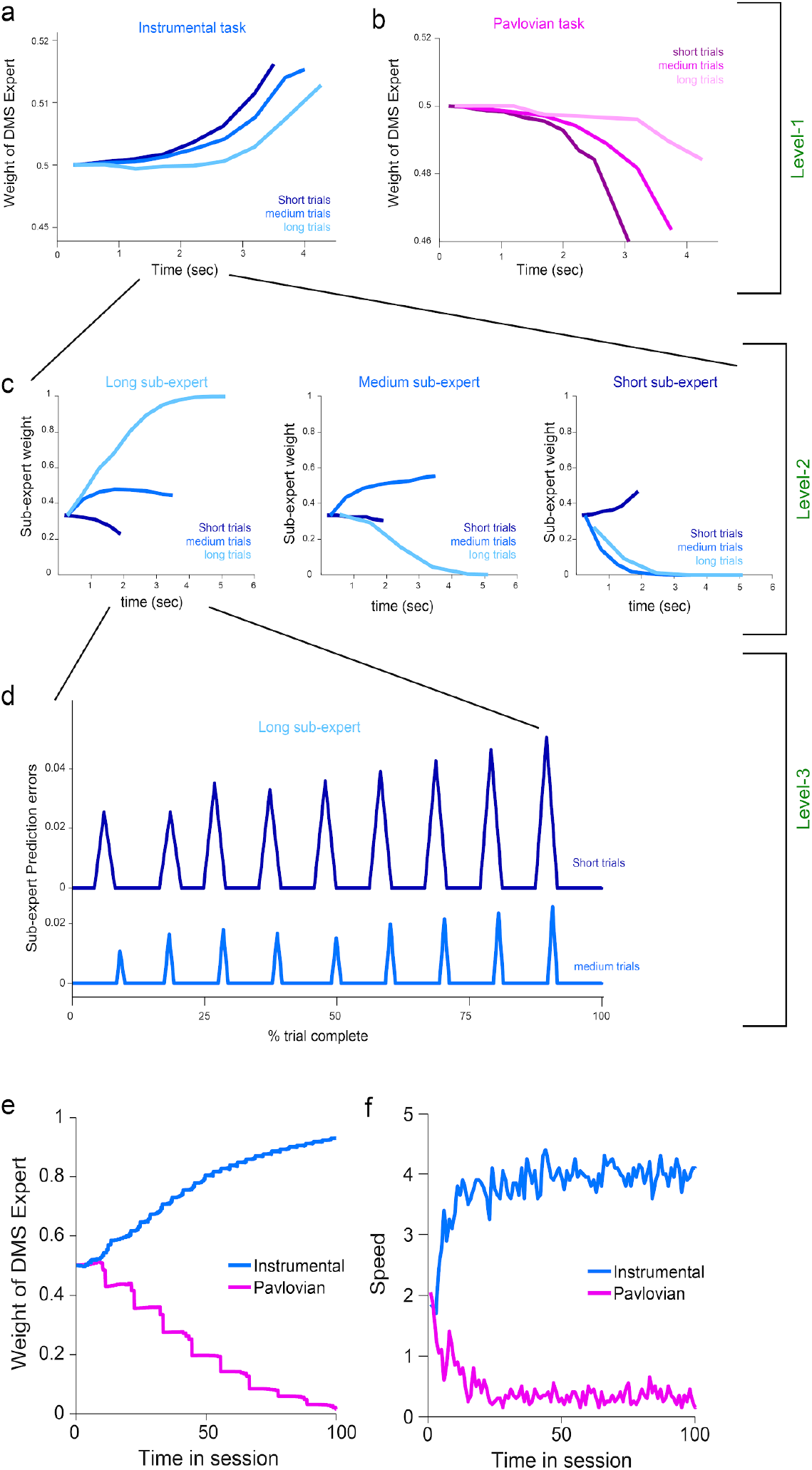
Within-trial dynamics of model variables at all three levels under different task conditions. **a,b** Positive and negative accumulation of distance expert (level-1, equivalent to DMS) weights under (a) instrumental and (b) Pavlovian task condition, for short, medium and long trial types. Each trace is the average dynamics on the very first trial, averaged for 10 simulation sessions. Similar dynamics accumulate across trials within a session when the task is repeated (not shown). Note that the ramp shape is convex in the first trial but concave for later trials. **c,** Within the distance expert, sub-experts (level-2) specialize on distinct contingencies and the weights ramp accordingly depending on task conditions. **d,** RPEs within a sub-expert in which tone transitions occur at unexpected times/distances (RPEs are zero for sub-experts that perfectly predict the current contingency; not shown). Note the larger magnitude RPEs for short compared to longer trials, as seen empirically (Fig. 5). Escalation of RPEs across the trial is due to temporal discounting. Similar to the empirical data, the impact of larger RPEs on short distances is more evident later in the trial. **e,** Example evolution of DMS (distance) expert weights across a session. Weights accumulate across trials to provide evidence the agent is in control. **f,** Model velocities (averaged across simulations) recapitulate increase in running in instrumental compared to pavlovian sessions. The model selects speeds in proportion to inferred responsibility of the distance expert.

**Supplementary Fig. 8:**
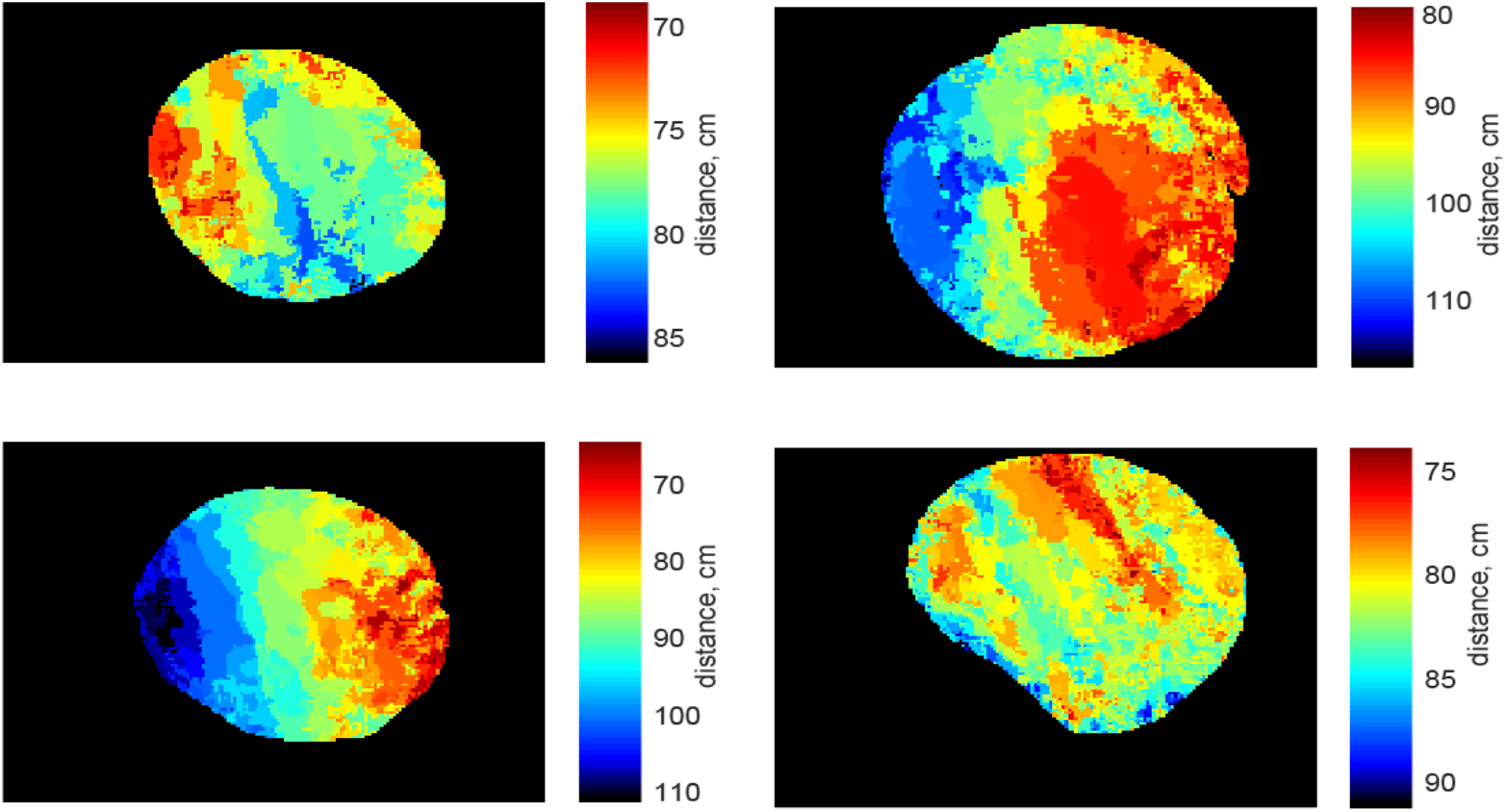
Additional example session of striatal subregions that have preferred distance contingency. The trial-by-trial ramp slope during anticipation epoch was distinctly modulated for different striatal subregions. Color maps are in same format as Fig. 4n.

**Supplementary Fig. 9:**
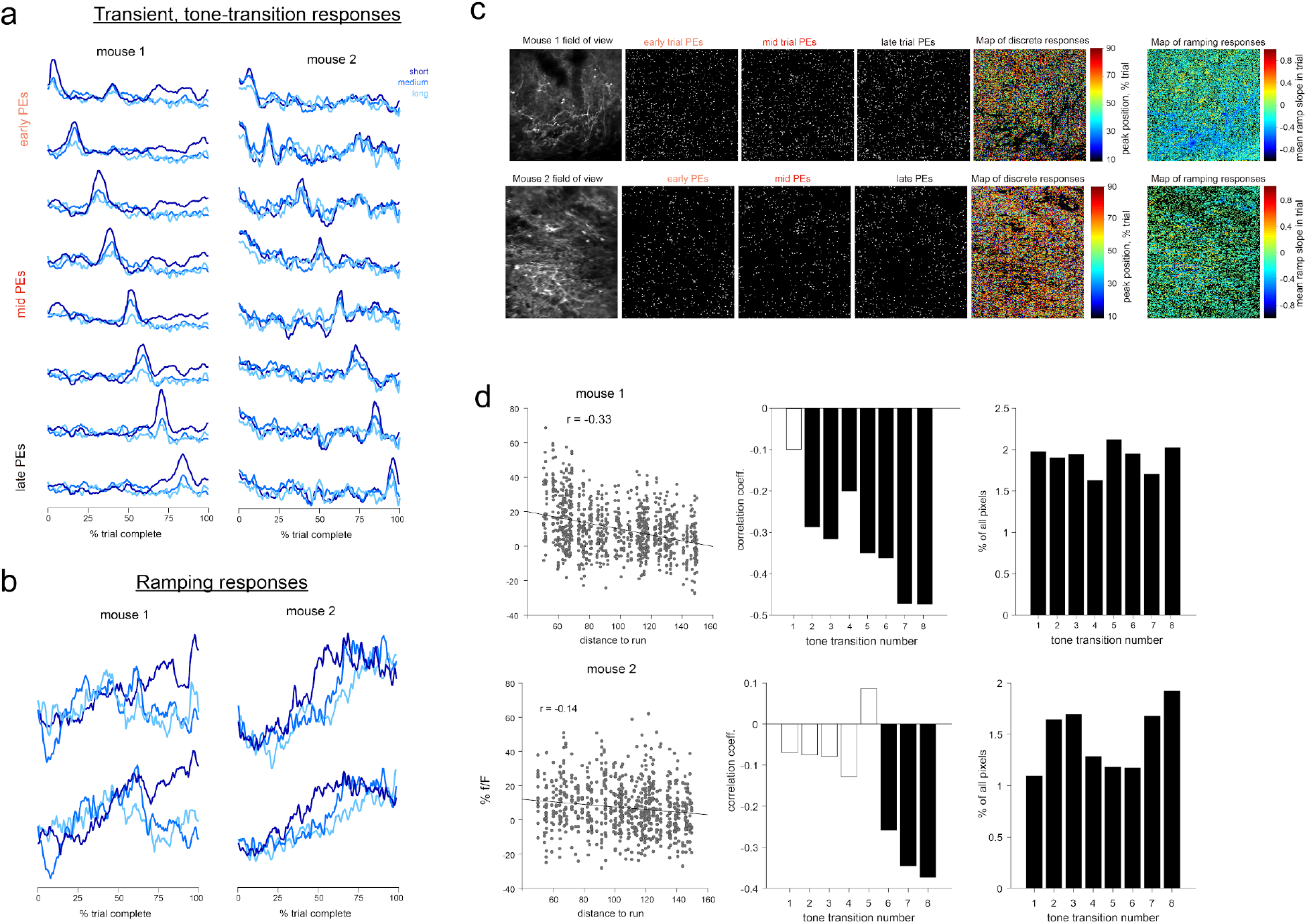
Discrete and ramp-like responses in dopamine axon segments. **a,** Discrete, tone responses in two mice. Format is as in Fig 5. **b,** Example ramping patterns in the two animals. **c,** Examination of the anatomical distribution of pixels that exhibit tone-transition tuning. *Top* row summarizes data from mouse-1, and *bottom* is for the second mouse. Leftmost panels show the mean projection of field of view, and the next three panels show the individual pixels that display discrete responses early (first transition), mid-trial (5th transition) and late-trial (last-tone transition) responses. Moreover, the anatomical organization of these pixels are intermingled. Fifth panel shows the anatomical position of all phasically responding pixels color coded for which transition they respond to. Finally, Last panel on the right shows the anatomical distribution and trial-averaged ramp slopes of pixels within the 2-photon field of view that exhibit sustained upward or downward activity during the anticipatory epoch. Same format for mouse-2 at *bottom* **d,** Quantification of how peak response at tone-transition is affected by distance to needed to run on current trial. Our simulations predict (see Supplementary Fig. 7) that shorter trials will elicit larger PEs. We found a significantly negative correlation overall in both mice (p<0.001, *left*), but the influence of distance was more prominent for later tones (*middle*, filled bar are have p<0.05) as in the model (Supplementary fig 7). Right panel shows that similar fraction of pixels that were responsive to each tone transition.

**Supplementary Video 1:** Example recording session demonstrating the activity pattern of dopamine axons in the dorsal striatum. Video playback is 1X.

**Supplementary Video 2:** Video illustrating extraction of flow trajectories on a frame by frame basis. Video playback is slowed down 0.25X.

**Supplementary Video 3:** Reward response in Instrumental task. Clock at top left displays time relative to reward. Video playback is 1X.

**Supplementary Video 4:** Reward response in Pavlovian task. Clock at top left displays time relative to reward. Video playback is 1X.

**Supplementary Video 5:** Progressively organized and continuous reward response to unpredicted reward deliver in naive mice (*top*), or animals that have received training in pavlovian sessions for 3 weeks. Clock at top left displays time relative to reward. Video playback is slowed down 0.5X.

## Notes

#### Summary of Updates

Additional clarifications, typos, and a panel in SFig7. Also higher resolution images for all figures.

